# Intramembrane protease RHBDL4 cleaves oligosaccharyltransferase subunits to target them for ER-associated degradation

**DOI:** 10.1101/850776

**Authors:** Julia D. Knopf, Nina Landscheidt, Cassandra L. Pegg, Benjamin L. Schulz, Nathalie Kühnle, Chao-Wei Chao, Simon Huck, Marius K. Lemberg

## Abstract

The Endoplasmic Reticulum (ER)-resident intramembrane rhomboid protease RHBDL4 generates metastable protein fragments and together with the ER-associated degradation (ERAD) machinery provides a clearance mechanism for aberrant and surplus proteins. However, the endogenous substrate spectrum and with that the role of RHBDL4 in physiological ERAD is mainly unknown. Here, we use a substrate trapping approach in combination with quantitative proteomics to identify physiological RHBDL4 substrates. This revealed oligosacharyltransferase (OST) complex subunits such as the catalytic active subunit STT3A as substrates for the RHBDL4-dependent ERAD pathway. RHBDL4-catalyzed cleavage inactivates OST subunits by triggering dislocation into the cytoplasm and subsequent proteasomal degradation. Thereby, RHBDL4 controls the abundance and activity of OST, suggesting a novel link between the ERAD machinery and glycosylation tuning.

## Introduction

Control of the cellular proteome is an essential process in every cell. In order to grow, divide and survive under changing conditions, cells need to finely tune the amount of each of their proteins by regulating synthesis, modification and degradation. This occurs both at the transcriptional and translational level by affecting synthesis and degradation rates of both mRNA and proteins to tightly control the total amount of any protein (Vogel and Marcotte, 2012). The later in the biosynthesis pathway regulation takes place, the faster it allows the cell to respond to current demands. Moreover, a dynamic protein homeostasis (proteostasis) network ensures that damaged cellular proteins are efficiently removed and that protein complexes assemble in the correct stoichiometry (Juszkiewicz and Hegde, 2018). When not corrected by the protein degradation machinery, imbalance of protein stoichiometry generates a major fitness defect by leading to toxic protein species and protein aggregation (Brennan et al., 2019). The endoplasmic reticulum (ER) is the major site for protein synthesis, maturation and quality control for all proteins within the secretory pathway (Ellgaard et al., 2016). Here, post-translational control of protein abundance is mediated either by proteasomal degradation or the lysosomal system. The latter mainly targets terminally misfolded proteins and large substrates whose degradation occurs via selected autophagy or direct ER-to-lysosome trafficking (Fregno and Molinari, 2019). However, the major clearance pathway for misfolded proteins relies on the ER associated degradation (ERAD) pathway that targets proteins for degradation by the cytoplasmic proteasome (Christianson and Ye, 2014; Mehrtash and Hochstrasser, 2018; Ruggiano et al., 2014).

ERAD is commonly defined by its role in eliminating aberrant proteins that could cause harm to the cell. Depending on the location in the protein of the lesion or the signal for regulated degradation (degron), ERAD substrates are classified into three main categories: ERAD-L (lumen), ERAD-M (membrane) and ERAD-C (cytosol). In yeast, three main E3 ubiquitin ligases target proteins from the ER for proteasomal degradation: Hrd1 is known to recognize ERAD-L and most ERAD-M substrates whereas ERAD-C substrates are handled by Doa10 (Carvalho et al., 2006; Vashist and Ng, 2004). At the inner nuclear membrane, a subarea of the ER, the Asi complex targets mislocalized membrane proteins for proteasomal degradation (Foresti et al., 2014; Khmelinskii et al., 2014). In mammals, an even more complex proteostasis network including several additional ERAD E3 ligases and auxiliary factors selectively target ER proteins into several parallel degradation routes (Christianson and Ye, 2014; Mehrtash and Hochstrasser, 2018). For ERAD-M substrates much remains unknown about the structural features that target proteins for degradation. In some cases, the overall conformation of a transmembrane (TM) region rather than a sequence feature is recognized (Gardner and Hampton, 1999; Sato et al., 2009). On a sequence level, one common feature in TM domains that targets proteins for ERAD is the presentation of charged amino acids within the TM domain that would normally be masked by a binding partner, as has been shown for the unassembled T cell receptor α-chain (Bonifacino et al., 1990).

Besides degradation of misfolded proteins and subunits of multiprotein complexes present in excess of their proper stoichiometry, ERAD plays a regulatory role by degrading functional proteins in response to changing demands of the cell. This has been shown for several substrates including immune receptors (van den Boomen et al., 2014; Wiertz et al., 1996) or the degradation step in negative feedback loops regulating receptors, metabolite transporters (Adle et al., 2009; Avci et al., 2014), ion transporters (Pearce et al., 2007) or enzymes involved in lipid metabolism (Foresti et al., 2013; Sever et al., 2003; Suzuki et al., 2012). The low number of proteins described to date whose abundance is regulated by ERAD makes it difficult to characterize the machinery involved. Unlike classical ERAD substrates, these conditional substrates all have in common a lack of an obvious folding defect, meaning that they therefore potentially require a different combination of ER quality control and ERAD factors than conventional pathways (Mehrtash and Hochstrasser, 2018). Previous work has shown that ER-localized intramembrane proteases can serve as key decision makers that direct correctly folded proteins towards the ERAD pathway. Signal peptide peptidase (SPP) and its yeast orthologue Ypf1 have been suggested to interact with ERAD factors to target specific proteins for degradation (Avci et al., 2014; Boname et al., 2014; Chen et al., 2014).

Intramembrane proteases cleave peptide bonds within cellular membranes, a hydrophobic environment that is only rarely exposed to water (Sun et al., 2016). In humans, 15 different intramembrane proteases have been identified that fall into different mechanistic families: metalloproteases, GxGD aspartyl proteases, the glutamyl protease Rce1, and rhomboid serine proteases (Kühnle et al., 2019b). In mammals, 5 rhomboid proteases exist with RHBDL4 (also known by its gene name *Rhbdd1*) being the only one localizing to the ER (Fleig et al., 2012; Wunderle et al., 2016). RHBDL4 has been shown to be upregulated in the ER unfolded protein response (UPR) and to cleave unstable membrane proteins (Fleig et al., 2012) as well as aggregation-prone soluble ERAD-L substrates (Kühnle et al., 2019a). Despite this broad substrate spectrum, a common theme is that RHBDL4-catalyzed cleavage generates fragments that are targeted for proteasomal degradation by ERAD machinery. For membrane proteins, RHBDL4 uses a bipartite substrate recognition mechanism. First, it recognizes unstable or positively charged TM domains such as presented by the orphan α-subunit of pre-T cell receptor (Fleig et al., 2012; Recinto et al., 2018). Second, RHBDL4 recognizes substrates through a conserved ubiquitin-interacting motif (UIM) at the cytosolic C-terminal tail of RHBDL4 thereby linking it to canonical ERAD E3 ubiquitin ligases such as gp78 (Fleig et al., 2012). Besides this involvement in ERAD, RHBDL4 has been implicated in sterol regulated cleavage of the amyloid precursor protein (APP) (Paschkowsky et al., 2018) as well as directly or indirectly tuning secretion dynamics (Wunderle et al., 2016). However, so far, no comprehensive study on the substrate spectrum of RHBDL4 exists and we therefore set out to pursue an affinity-enrichment-based proteomic approach using the catalytically inactive mutant of RHBDL4 as a bait to identify unambiguously RHBDL4 substrates.

We identified several subunits of the oligosacharyltransferase (OST) complex as substrates for the RHBDL4-dependent ERAD pathway. The OST complex catalyzes the transfer of a preassembled oligosaccharide from a lipid-linked oligosaccharide donor onto the asparagine residue of glycosylation acceptor sites known as sequons (N-X-S/T; X≠P) in newly synthesized proteins (Cherepanova et al., 2016). Insects, vertebrates and plants assemble two OST complexes that are composed of a complex-specific catalytic subunit (STT3A or STT3B), a set of shared subunits (RPN1, RPN2, DDOST, DAD1, OST4 and TMEM258 in mammalian cells) as well as complex-specific accessory subunits that only assemble with STT3A (DC2 and KCP2) or STT3B (MagT1 or TUSC3) (Kelleher et al., 2003; Roboti and High, 2012; Shibatani et al., 2005). STT3A catalyzes co-translational glycosylation and is found in complex with the Sec61 translocon channel. A major cellular role for the STT3B complex is to maximize sequon occupancy by glycosylating acceptor sites that are skipped by the STT3A complex (Cherepanova et al., 2019; Shrimal et al., 2015). In addition, it has been shown that post-translocational glycosylation of cryptic sequons by STT3B, that in the native fold are not modified, targets proteins to the glycan-dependent branch of the ERAD pathway (Sato et al., 2012). Little is known about regulation of OST abundance. While several studies have shown that the ER undergoes major structural and functional changes during the UPR leading to decreased protein translation and import and increased ER export (Hetz and Papa, 2018; Walter and Ron, 2011), it is unclear whether glycosylation activity is also modulated. RPN1 and RPN2, which in addition to their role in the OST complex have been proposed to act as molecular chaperones (Qin et al., 2012), are UPR target genes. On the other hand, the catalytic subunits STT3A and STT3B appear not to be affected by the UPR (Dombroski et al., 2010). While different subunits are not strictly co-regulated, proper OST complex stoichiometry has to be maintained. In yeast, it has been shown that ERAD compensates for disturbed homeostasis, such as observed in heterozygous diploid cells, by degrading excess subunits (Mueller et al., 2015). Recent work showed that OST subunits failing to assemble in functional complexes in yeast are recognized by the inner nuclear membrane Asi E3 Ligase (Natarajan et al., 2019). Yet, how these natively folded proteins are discriminated from stable inner nuclear membrane proteins and whether a related mechanism controlling OST assembly exits in higher organisms, which seem not to have Asi homologues, remain important questions. By following a substrate proteomics approach, we show that the ERAD rhomboid protease RHBDL4 cleaves several OST subunits leading to their dislocation into the cytoplasm and degradation by the proteasome. We propose that cleavage by RHBDL4 serves as a potent and irreversible degradation signal to drive turnover of otherwise stable OST complexes in order to adapt glycosylation activity to functional need.

## Results

### Substrate screen identifies OST complex subunits as RHBDL4 substrates

In order to identify endogenous substrates of RHBDL4 we performed a screen in Hek293T cells using stable isotopic labeling in cell culture (SILAC)-based quantitative proteomics. We have previously shown that the catalytically inactive RHBDL4 serine-144-alanine (SA) mutant can bind but not cleave cognate substrates and that this interaction is reduced after mutating the UIM of RHBDL4 (Fleig et al., 2012). We therefore reasoned that the SA mutant can be used as a substrate trap in order to unambiguously identify endogenous RHBDL4 substrates. To compare substrate interaction with all three RHBDL4 variants, we generated doxycycline-inducible Hek293 T-REx cells stably expressing C-terminally GFP-tagged RHBDL4 wild-type (wt), the SA mutant or the RHBDL4-SA-UIM double mutant (SAUM) (Fig. S1A) and performed a triple SILAC experiment with RHBDL4 wt in “light”, SAUM in “medium” and SA in “heavy”. One-step affinity purification of the GFP-fusion proteins was performed from detergent-solubilized microsomal membranes of equal amounts of cells mixed as outlined in Fig. 1A. Mass spectrometry (MS) analysis identified 273 proteins quantified in both replicates of which 47 were integral ER membrane proteins (Table S1 and 2). Of these, 32 proteins were enriched with a mean equal or greater 2.0 in the precipitate of RHBDL4-SA when compared to RHBDL4 wt (Table 1). 10 of the substrate candidates are polytopic membrane proteins, which is therefore the most enriched protein topology. All identified proteins showed a reduced enrichment with RHBDL4-SAUM, indicating that their binding is enhanced upon ubiquitination (Table 1). Among the identified substrate candidates were four proteins, RPN1, STT3A, DDOST and RPN2, that all belonged to the OST complex. Trapping of both RPN1 and STT3A was independently validated, though only a minor fraction of the total cellular pool was recovered (Fig. S1B). Taken together, these results indicate that RHBDL4 interacts with a subpopulation of these endogenous ER-resident proteins to trigger their degradation. Next, we asked whether RHBDL4 directly proteolytically processes OST subunits by performing a cell-based rhomboid gain-of-function cleavage assay (Adrain et al., 2011; Fleig et al., 2012). In order to detect potential low abundant cleavage fragments, we generated expression constructs with an N-terminal triple FLAG-tag (Fig. 1B). Overexpression of RHBDL4 wt but not of the SA mutant generated several N-terminal cleavage fragments for both RPN1 and STT3A, indicating a direct role for rhomboid-catalyzed cleavage (Fig. 1C and D). However, the level of full-length substrates did not significantly decrease, indicating that only a portion is affected and for cleavage additional steps such as ubiquitination may be required as observed before for RHBDL4-catalyzed processing of model ERAD substrates (Fleig et al., 2012). Interestingly, RHBDL4-generated RPN1 cleavage fragments ranged from 23 kDa to 55 kDa (Fig. 1C). Considering RPN1 topology (Fig. 1B), this result suggests that, RHBDL4 cleaves RPN1 at several positions in the luminal domain and once in the TM segment. For STT3A, two internal cleavage fragments are generated by RHBDL4 of 35 kDa and 13 kDa, respectively (Fig. 1D, see below for further characterization of cleavage sites). Inhibition of the proteasome with MG132 increased steady-state levels of the RPN1 and STT3A fragments (Fig. 1C and D), indicating that they become dislocated into the cytoplasm for proteasomal degradation, as has been previously observed (Fleig et al., 2012). This is also consistent with previous reports on proteasomal degradation of excess OST subunits in yeast (Mueller et al., 2015; Natarajan et al., 2019). We also validated RPN2 and DDOST as RHBDL4 substrates and could show that they are prone to RHBDL4-catalyzed cleavage (Fig. S1C-E). STT3A is the catalytic subunit of the dominant OST complex that mediates co-translational N-glycosylation of most sequons, while STT3B-containing complexes are mainly required for efficient post-translational glycosylation of sites that have been skipped by STT3A (Shrimal et al., 2015). Interestingly, STT3B was also identified in one replicate of our proteomics screen and enriched by 30 % in the RHBDL4-SA precipitate when compared to RHBDL4 wt (Table S1 and 2). We therefore asked whether STT3B is indeed cleaved by RHBDL4, as both STT3 proteins have a sequence similarity of 63.5 % with the main difference being a 51-amino acid extension at the N-terminus of STT3B (Fig. 1B). In our cell-based gain-of-function assay ectopically expressed STT3B was cleaved, resulting in a 16-kDa cleavage fragment that, considering the sequence variation at the N-terminus, corresponds to the 13-kDa STT3A-fragment (Fig. 1E). Our screen also identified another protein linked to glycosylation as an RHBDL4 substrate, the type II membrane protein β-1,4-glucuronyltransferase also known as B4GAT1 (Fig. S1C and F). This cleavage, which is unexpected because rhomboids initially had been described to cleave only type I membrane proteins (Dumax-Vorzet et al., 2013), is predicted to take place in the luminal ectodomain of B4GAT1. Taken together with a previous report that *Drosophila* Rhomboid-1 can cleave the type II membrane protein Star (Tsruya et al., 2007), this result suggest that rhomboids have an even broader substrate spectrum than anticipated.

**Table 1.** Result of MS/MS based screening for RHBDL4 substrates.

| **Protein ID** | **Gene name** | **Mean M/L** | **Mean H/L** | **Unique Peptides** | **TMDs** |
| --- | --- | --- | --- | --- | --- |
| O15260 | SURF4 | 1.40 | 2.88 | 9 | 5 |
| O15269 | SPTLC1 | 1.31 | 2.80 | 7 | 1 |
| Q7Z3C6 | ATG9A | 1.07 | 2.77 | 15 | 6 |
| P37268 | FDFT1 | 1.36 | 2.70 | 15 | 1 |
| P50402 | EMD | 1.65 | 2.68 | 12 | 1 |
| Q14165 | MLEC | 1.12 | 2.62 | 5 | 1 |
| O95573 | ACSL3 | 1.34 | 2.62 | 22 | 1 |
| P04843 | RPN1 | 1.22 | 2.60 | 41 | 1 |
| Q53GQ0 | HSD17B12 | 1.24 | 2.59 | 9 | 3 |
| Q9UBM7 | DHCR7 | 1.52 | 2.48 | 8 | 7 |
| Q9UBV2 | SEL1L | 1.24 | 2.45 | 15 | 1 |
| P46977 | STT3A | 1.19 | 2.43 | 16 | 13 |
| Q14318 | FKBP8 | 1.05 | 2.40 | 11 | 1 |
| P39656 | DDOST | 1.22 | 2.37 | 16 | 1 |
| O95864 | FADS2 | 1.50 | 2.36 | 7 | 4 |
| Q99523 | SORT1 | 1.48 | 2.36 | 12 | 1 |
| Q9Y5M8 | SRPRB | 1.12 | 2.33 | 7 | 1 |
| Q969M3 | YIPF5 | 1.31 | 2.32 | 4 | 5 |
| Q07065 | CKAP4 | 1.28 | 2.27 | 20 | 1 |
| Q9BTV4 | TMEM43 | 1.22 | 2.26 | 22 | 4 |
| Q15043 | SLC39A14 | 1.48 | 2.24 | 8 | 6 |
| Q96A33 | CCDC47 | 1.33 | 2.23 | 14 | 1 |
| P04844 | RPN2 | 1.15 | 2.22 | 8 | 3 |
| Q9NZ01 | TECR | 1.20 | 2.19 | 17 | 3 |
| P16615 | ATP2A2 | 1.23 | 2.18 | 83 | 10 |
| P05023 | ATP1A1 | 1.36 | 2.16 | 56 | 10 |
| O00264 | PGRMC1 | 0.95 | 2.15 | 9 | 1 |
| P27824 | CANX | 1.13 | 2.15 | 38 | 1 |
| Q96JJ7 | TMX3 | 1.46 | 2.14 | 12 | 1 |
| P43307 | SSR1 | 1.20 | 2.12 | 4 | 1 |
| Q6ZMG9 | CERS6 | 1.21 | 2.09 | 4 | 5 |
| P20020 | ATP2B1 | 1.36 | 2.00 | 19 | 10 |

**Fig. 1.**
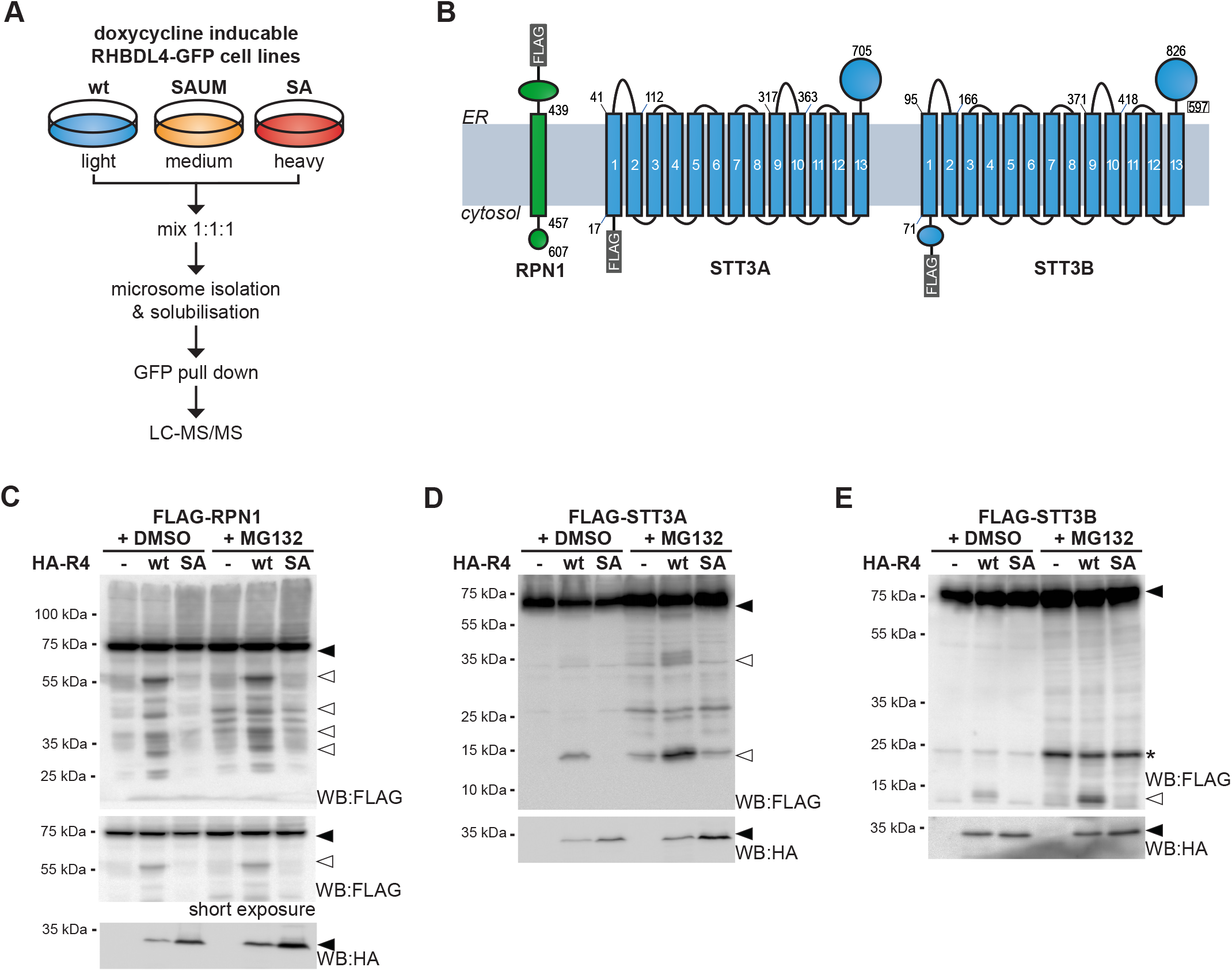
Substrate trapping identifies OST complex subunits as RHBDL4 substrates. (A) Experimental outline of SILAC-based mass spectrometry analysis of RHBDL4 ‘trappome’ from Triton X-100-solubilized Hek293 T-REx-RHBDL4-GFP cells. (B) Outline of used N-terminally FLAG-tagged constructs for FLAG-RPN1, FLAG-STT3A and FLAG-STT3B. Numbers indicate positions of critical TM domain boundaries. (C) Hek293T cells were co-transfected with FLAG-RPN1 (filled triangle) and either an empty vector (−), HA-tagged RHBDL4 (HA-R4) wt, or RHBDL4-SA. RHBDL4 generates N-terminal cleavage fragments (open triangle) that are degraded by the proteasome as shown by increased steady state level upon MG132 treatment (2 μM) compared to vehicle control (DMSO) in the displayed short exposure. (D) Cleavage assay as shown in (B) but with FLAG-STT3A as substrate. (E) Cleavage assay as shown in (B) but with FLAG-STT3B-FLAG as substrate.

### RHBDL4 is required for OST complex homeostasis

In the cell-based cleavage assay above, orphan subunits of an oligomeric complex were generated by ectopically expressing them in excess. We wondered whether RHBDL4 also recognizes orphan subunits expressed at physiological level if they lack their respective complex partners. It has been shown previously that knockdown of the OST subunit OST4 leads to reduced steady state levels of the other complex partners, but the underlying molecular mechanism of these reduced levels have not yet been resolved (Dumax-Vorzet et al., 2013). We therefore asked whether RHBDL4 plays a role in degradation of orphan OST subunits when expressed at endogenous level. Yet we needed to overcome the challenge of antibodies being often not sensitive enough to detect subtle changes in protein expression. We therefore endogenously tagged both STT3 subunits at their C-terminus with a triple FLAG-tag using CRISPR/Cas12-mediated gene editing (Fueller et al., 2019), allowing for sensitive western blot detection (Fig. S2A-C). Indeed, as had been previously observed, knockdown of OST4 leads to a subtle reduction of the steady-state level of STT3A (Fig. 2A and S2D) (Dumax-Vorzet et al., 2013). In contrast, double knockdown of OST4 and RHBDL4 shows a significant increase of STT3A levels partially rescuing this destabilizing effect. This result shows that RHBDL4-dependent ERAD contributes to removal of endogenous orphan OST complex subunits, thereby controlling the stoichiometry of complex assembly and clearing the ER from unassembled complex subunits. A similar influence could not be observed for STT3B (Fig. 2B and S2D). Whereas previously no effect of OST4 depletion on STT3B has been observed (Dumax-Vorzet et al., 2013), in our hands short term depletion of OST4 resulted in a ~10% decrease of STT3B steady-state levels. However, in contrast to STT3A complex destabilization this could not be modulated by RHBDL4 depletion, indicating that under these conditions a different ERAD pathway acts on STT3B.

**Fig. 2.**
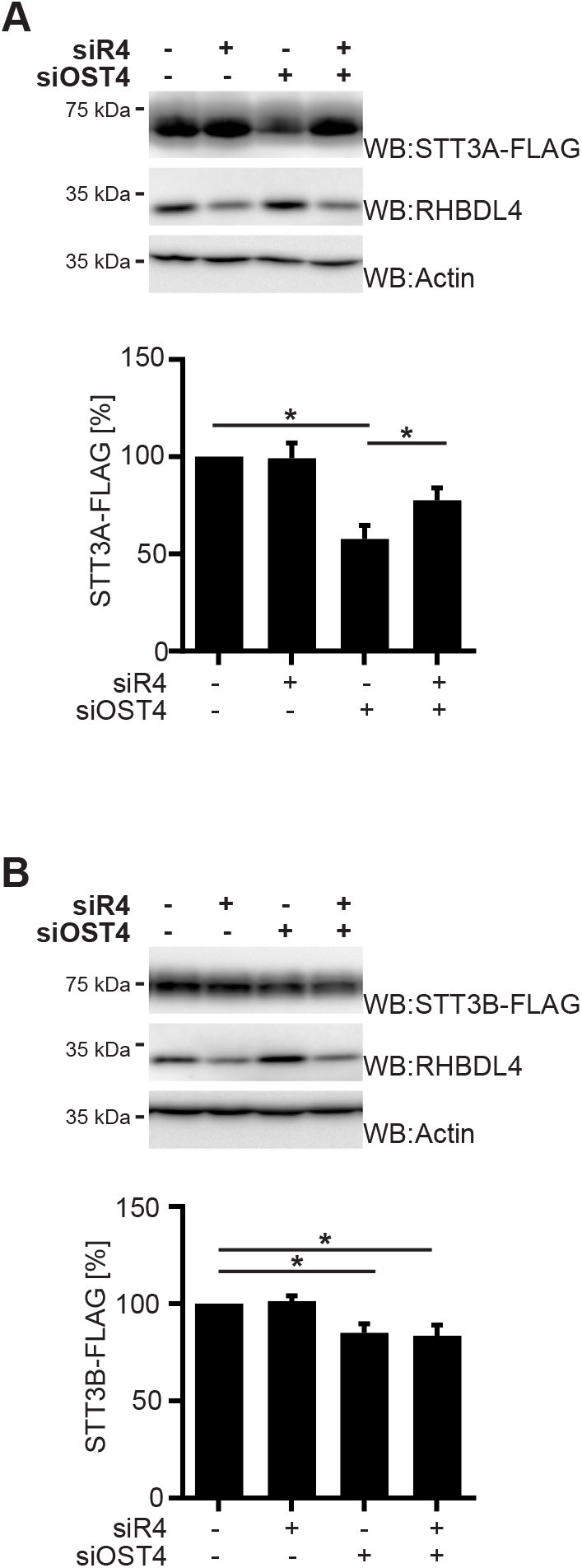
RHBDL4 is required for OST complex homeostasis. (A) Hek293T cells expressing C-terminally tagged STT3A (STT3A-FLAG) from the endogenous locus were transfected with siRNA targeting either RHBDL4 (siR4), OST4 (siOST4), a combination of both or control siRNA (sictrl) only. Actin serves as loading control. Lower panel: quantification of STT3A-FLAG expression level relative to sictrl treated cells (means ± SEM, n=5; *p < 0.05, one-way ANOVA). WB, western blot. (B) Assay as in (a) but with Hek293T cells expressing STT3B-FLAG from the endogenous locus. Lower panel: quantification of STT3B-FLAG expression level relative to sictrl treated cells (means ± SEM, n=6, *p < 0.05, one-way ANOVA).

### RHBDL4 cleaves STT3A at two distinct luminal loops

Our proteomics result suggests that RHBDL4 cleavage has a major impact on degradation of polytopic membrane proteins. Our preceding work with mutated model ERAD substrates of this topology suggested that cleavage can occur in luminal loops and TM segments (Fleig et al., 2012). Yet, what defines a RHBDL4 cleavage site is, due to the lack of characterized endogenous substrate cleavages, not clear. Hence, we aimed to further characterize STT3A cleavage, which occurred in at least two defined regions. First, we mapped cleavage sites by comparing RHBDL4-generated fragments with truncated versions of STT3A as reference peptides (Fig. S3A and B). The 13-kDa fragment detected in presence of MG132 could be narrowed down to loop 1 (L1) whereas the 35-kDa fragment results from a cleavage event in loop 9 (L9), which both are located within the ER lumen (Fig. S3D). Cleavage site requirements for rhomboid protease substrates still remain elusive. For bacterial rhomboid proteases small side chains at the scissile peptide bond had been shown to at least partially determine cleavage specificity (Strisovsky et al., 2009). In the L1 region corresponding to the 13-kDa fragment there was only one residue matching this requirement, namely glycine-64. Mutating this position to phenylanalyine (G64F) reduced the amount of this cleavage catalyzed by ectopically expressed RHBDL4 by about 50% (Fig. 3A). Likewise, processing within the L1 region was reduced in the control experiment, indicating that endogenous RHBDL4 also recognizes glycine-64 as a cleavage site. Interestingly, this glycine is conserved in STT3B (G118) and, considering the approximate 50 amino acid cytoplasmic N-terminal extension of STT3B, this result suggests that the 16-kD fragment of STT3B is also generated by cleavage at this position (Fig. 1E). Cleavage in L9 could be narrowed down to near position 350, close to two serine residues at 348 and 350. Mutating serine-348 to phenylanalyine (S348F), but not serine-350 (S350F), reduced the amount of the 35-kDa cleavage fragment generated upon RHBDL4 overexpression by about 40% (Fig. 3A and S3C). Taken together, these results suggest that STT3A can be cleaved at two alternative luminal sites, with L1 and L9 being the only larger luminal protrusions present in the structure of STT3 (Fig. 3B and S3D). Interestingly, the cleavage site in L9 is located at the binding interface of STT3B with its two paralog-specific subunits MagT1 and TUSC3, which in STT3A-type OST complexes is occupied by DC2 (Braunger et al., 2018; Ramirez et al., 2019). However, in contrast to MagT1 or TUSC3, DC2 does not possess any luminal segments that might ‘mask’ the cleavage site and probably prevent processing at this position, explaining why this cleavage site is specific for STT3A. As the G64F and the S348F single and combined mutations did not completely abolish RHBDL4-catalyzed cleavage, the previously proposed rhomboid cleavage site motif (Strisovsky et al., 2009) and small side chains at the scissile peptide bond seem to be beneficial but not strictly required for RHBDL4 cleavage.

**Fig. 3.**
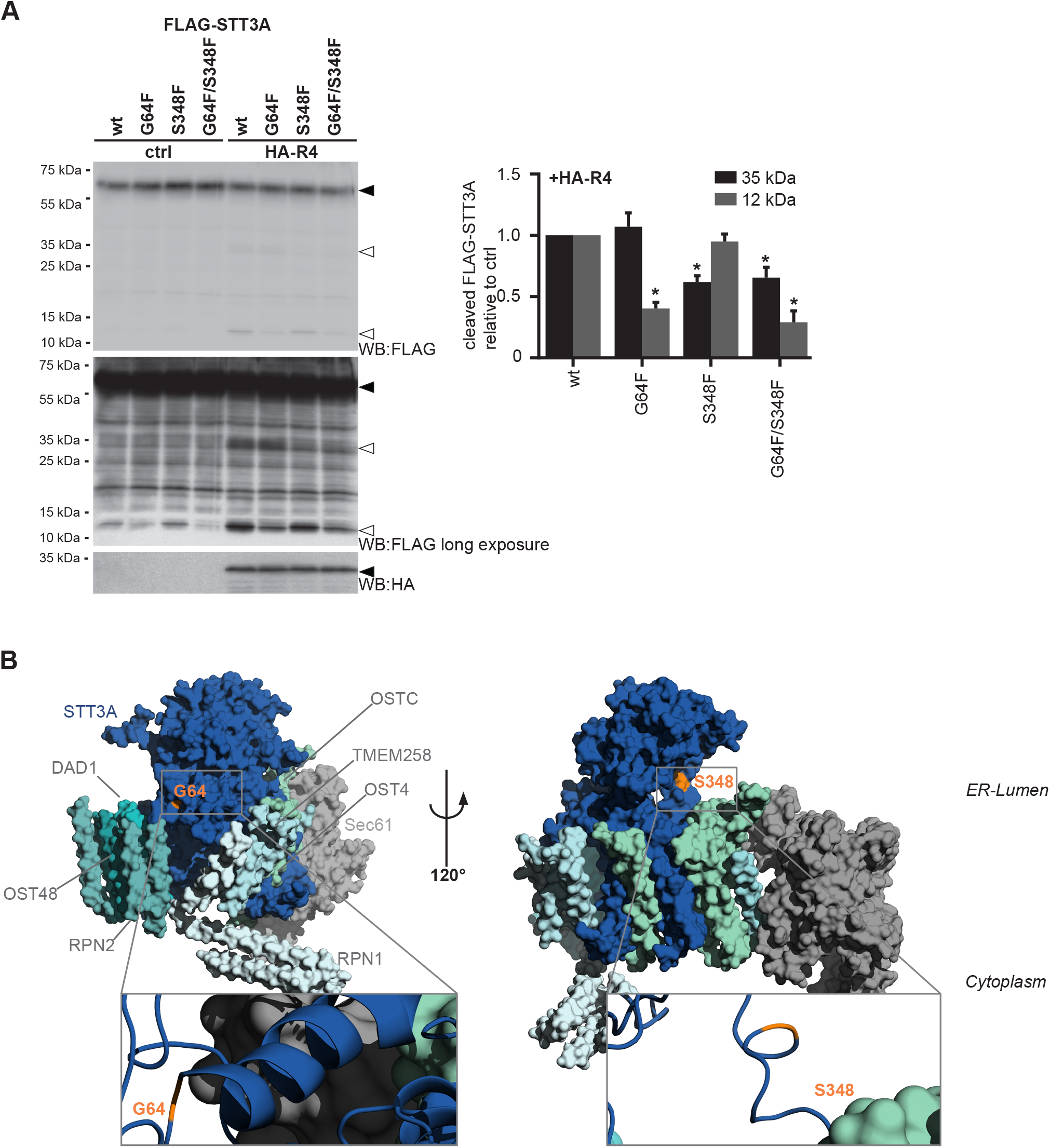
RHBDL4 cleaves endogenous STT3A in the ER lumen. (A) Hek293T cells were transfected with FLAG-STT3A wt, or with a G64F, S348F, or a G64F/S348 double mutant of FLAG-STT3A (filled triangle) together with HA-tagged RHBDL4 (HA-R4) constructs as indicated. Overexpression of the active rhomboid (wt) resulted in cleavage fragments at 35 kDa (L9) and 13 kDa (L1) (open triangles) observed significantly less for the SA mutant. In order to prevent clearance of cleavage fragments, cells have been treated with proteasome inhibitor MG132 (2 μM). Right panel, western blot (WB) quantification of respective cleavage fragment upon HA-R4 overexpression (means ± SEM, n=3; *p < 0.05, one-way ANOVA). Lower panel shows an oversaturated western blot. (B) Structure of STT3A (dark blue) in complex with other OST complex subunits and Sec61 (as indicated) taken from the coordinates from (Braunger et al., 2018) (PDB code 6FTI). Putative RHBDL4 cleavage sites at G64 (L1) and S348 (L9) are highlighted in orange.

### Endogenous STT3A is cleaved by RHDBL4

We next asked whether we can also monitor RHBDL4-dependent cleavage fragments if STT3A is expressed from its endogenous promotor. Facing the challenge of detecting low abundant cleavage fragments while only a subpopulation of the endogenous STT3A pool is affected by RHBDL4 cleavage, we took advantage of our endogenously tagged STT3A cell-line (Fig. S2A and B) enabling enrichment and sensitive western blot detection. We overexpressed active RHBDL4 and monitored for potential endogenous cleavage fragments (Fig. 4A), which we enriched by anti-FLAG immunoprecipitation. Indeed, in the presence of RHBDL4 wt, but not of the SA mutant, a weak FLAG-tagged fragment became visible running as a diffuse band at ~45 kDa (Fig. 4A). STT3A harbors three glycosylation sites (N537, N544, N548) in its large C-terminal luminal portion, explaining the diffuse running pattern of both full-length and cleaved STT3A. We therefore treated the samples with endoglycosidase H (Endo H), which shifts the cleavage fragment to 30 kDa (Fig. 4A). This suggests that it corresponds to the 35 kDa N-terminal fragment observed for N-terminally tagged STT3A (Fig. 1D). As the fragment was also visible in the vehicle transfected control, we wondered if this was due to endogenous RHBDL4 activity. Indeed, upon transient knockdown of RHBDL4 a significant reduction of the fragment could be observed (Fig. 4B). Ectopic expression of mouse RHBDL4 wt rescued cleavage of endogenously tagged STT3A (Fig. 4B). Taken together, these results show that a small fraction of endogenous STT3A can be cleaved by endogenous RHBDL4 leading to metastable fragments that become degraded by ERAD.

**Fig. 4.**
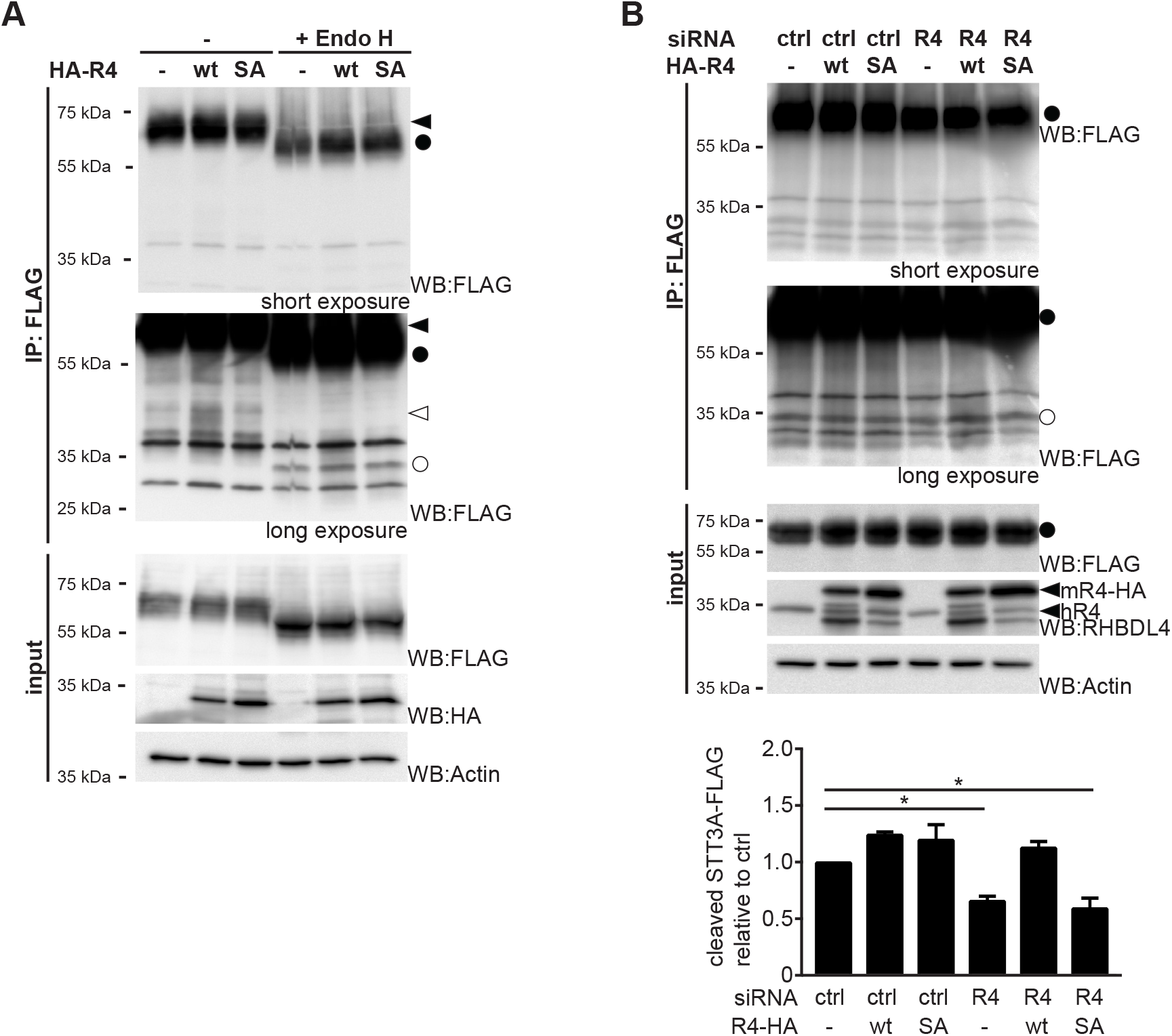
Endogenous STT3A is cleaved by RHBDL4. (A) Hek293T cells expressing C-terminally tagged STT3A (STT3A-FLAG) from the endogenous locus were transfected with either empty vector (−) or HA-tagged RHBDL4 (HA-R4) constructs as indicated. Full-length STT3A-FLAG (filled triangle) and its C-terminal cleavage fragments (open triangle) were isolated by anti-FLAG immunoprecipitation (IP). Treatment with Endo H prior western blotting (WB) removes N-linked glycans of STT3A and cleavage fragment (closed and open circle, respectively). In order to prevent clearance of cleavage fragments, cells have been treated with proteasome inhibitor MG132 (2 μM). Actin levels are shown as loading control. Long exposure shows an oversaturated western blot. (B) Hek293T-STT3A-FLAG cells have ben transfected either with control siRNA (ctrl) or siRNA targeting RHBDL4 (R4) and analyzed as in (A) by anti-FLAG immunoprectipitation and consequent EndoH digest. Ectopic expression of mouse HA-R4 rescues reduction of the STT3A-FLAG cleavage fragment (open circle). Actin levels are shown as loading control. Long exposure shows an oversaturated western blot. Actin levels are shown as loading control. Below, WB quantification of STT3A cleavage fragment relative to cells transfect with ctrl siRNA and empty plasmid (means ± SEM, n=3; *p < 0.05, one-way ANOVA).

### Two unique charges in STT3A facilitate RHBDL4-catalyzed cleavage

Previously, our work demonstrated that positive charges can serve as a TM degron for RHBDL4 mediated ERAD (Fleig et al., 2012). We wondered whether these features were also present in the TM domains of STT3A and STT3B. For STT3A, all TMs except TM helix 10 (H10), and for STT3B all except H5 and H9, exhibit at least one positively charged residue (Fig. S4A). Most of these charges either face other complex subunits or are located at the predicted membrane head group region facing either the cytosol or the ER lumen (Braunger et al., 2018; Wild et al., 2018). Interestingly, for STT3A 5 positive charges remain that can be clustered into two groups facing the plane of the membrane on different sides. Group one (I.) consists of three positive charges in H1 (K19), H2 (H133) and H5 (K197) (Fig. 5A and B). The second group (II.) consists of two positive charges: a histidine in TM segment H8 (H280) and an arginine in H9 (R300) (Fig. 5A and B). As all these 5 positively-charged amino acids are absent in STT3B and face facing the hydrophobic lipid acyl-chains of the membrane, we wondered whether they impact RHBDL4-catalyzed cleavage. We generated charge mutants of STT3A by replacing either single charged residues or all residues of one group with the respective residues of STT3B. Overall, this led to a reduced cleavage at glycine-64 but did not abolish cleavage completely (Fig. S4B) indicating that, at least if STT3A is expressed in excess, charged TM residues are not a strict requirement for cleavage. We therefore investigated whether transferring the STT3A-specific charges onto the STT3B paralogue can increase RHBDL4-dependent cleavage. We introduced one positive charge per group, L184H and Y331H, corresponding to STT3A H133 and H280, respectively (Fig. S4A). Indeed, this STT3B-L184H/Y331H double charge mutant, which on SDS-PAGE showed an unexpected accelerated electrophoretic mobility, exhibited increased RHBDL4 cleavage (Fig. 5C). Consistent with a beneficial role of positive charges in substrate recognition, the other polytopic RHBDL4 substrate RPN2 also has multiple positive TM charges (Fig. S4C). However, we note that several RHBDL4 substrates with a single TM domain, namely RPN1, DDOST, B4GAT1 (Fig. S4D) and APP (Paschkowsky et al., 2016), do not have any positive charges in their predicted TM segments. This indicates that, as with other ERAD substrate receptors (Christianson and Ye, 2014), RHBDL4 recognizes multiple substrate features.

**Fig. 5.**
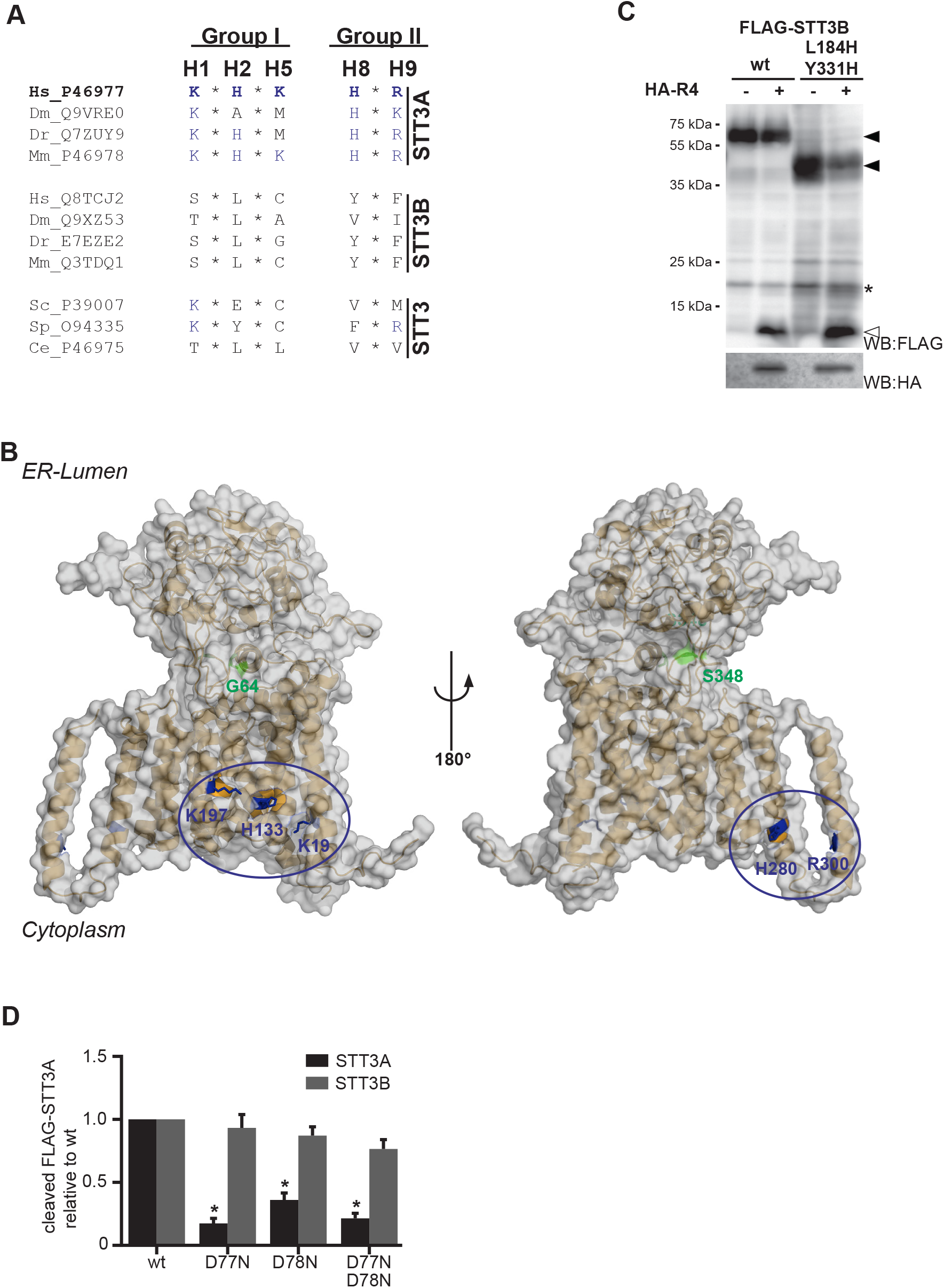
STT3A exhibits two unique charges missing in ancestral STT3 homologs. (A) Overview of paralogue specific charged TM residues found in human STT3A (Homo sapiens, Hs) but lacking in Hs STT3B and STT3B from mouse (Mus musculus, Mm), Danio rerio (Dr), Drosophila melanogaster (Dm) and Caenorhabditis elegans (Ce) as well in STT3 from Saccharomyces cerevisiae (Sc) and Schizosaccharomyces pombe (Sp). (B) Structure of human STT3A with paralogue specific positive charges highlighted in blue: group I (K19, H133, K197) and group II (H280, R300). RHBDL4 cleavage sites G64 (L1) and S348 (L9) in green (PDB code 6FTI). (C) FLAG-STT3B wt or STT3B-FLAG with positive charges in TM domain 3 and 8 (L184H/Y331H) have been transfected either with empty vector (-) or HA-tagged RHBDL4 (HA-R4) and cells have been treated with MG132 (2 μM). Of note, for FLAG-STT3B L184H/Y331H the six-fold amount of cDNA was transfected in order to allow comparison with the more stable FLAG-STT3B wt. WB, western blot. (D) Hek293T cells were transfected with FLAG-STT3A or FLAG-STT3B together with either HA-R4 wt, SA or mutants where RHBDL4-specific negative charged aspartate residues in TM H2 have been replaced by uncharged asparagine (D77N, D78N, D77N/D78N). RHBDL4-generated cleavage fragments (13-kDa for STT3A and 16-kDa for STT3B) have been quantified relative to the respective cleavage fragment generated by RHBDL4 wt. (means ± SEM, n=3; *p < 0.05, one-way ANOVA). See Fig. S4G for representative WB.

Strikingly, RHBDL4 has a conserved cluster of negatively-charged residues consisting of aspartate-77 and aspartate-78 that is missing from other rhomboid proteases (Fig. S4E). Based on the crystal structure of the *Escherichia coli* rhomboid protease GlpG (Wang et al., 2006), both these residues are predicted to be located at the luminal/membrane interface of TM segment H2 (Fig. S4F). Since for *E. coli* GlpG H2 and H5 have been implicated to act as a lateral active site gate (Cho et al., 2019), we asked whether the RHBDL4-specific negative charged residues contribute to substrate recruitment. To test this idea, we mutated both residues in the RHBDL4 H2 to structurally related but uncharged asparagine (D77N/D78N). Ectopically expression of both single mutants, and even more pronounced the double mutant, showed a reduced activity in cleaving STT3A (Fig. 5D and Fig. S4G). In contrast, the RHBDL4 H2 mutants showed robust activity against STT3B, demonstrating that the mutation does not compromise the overall rhomboid active site. Taken together with the facultative recognition of positive-charged substrate TM residues, this result suggests that a negatively-charged TM segment H2 assists recruitment of certain RHBDL4 substrates.

### RHBDL4 tunes ER glycosylation capacity

In order to study whether RHBDL4 plays a role beyond control of OST complex assembly, we performed cycloheximide chase experiments with the chromosomally tagged cell lines to study degradation rates of both STT3 subunits at endogenous levels and stoichiometry. Within 8 hours STT3A was degraded by about 25%, an effect that was completely abolished upon transient knock-down of RHBDL4 (Fig. 6A). In contrast, STT3B remained fully stable during the 8-hour chase period (Fig. 6B). This STT3A specific effect is consistent with our result observed for insufficient assembled OST complexes upon OST4 is knocked down (Fig. 2B). Next, we wondered whether OST degradation could be stimulated when the level of potential OST substrates becomes reduced as predicted to occur under chemically inducing UPR (Walter and Ron, 2011). Therefore, we induced ER stress by inhibiting the sarco/endoplasmic reticulum calcium-ATPase (SERCA)-mediated calcium uptake with cyclopiazonic acid (CPA) (Pirot et al., 2006) 24 hours prior to performing the chase experiment. In this treatment, degradation rate of STT3A was only mildly increased resulting in turnover of ~30% of STT3A after an 8-hour chase experiment (Fig. 6A). However, the otherwise stable STT3B was now also degraded by 30%, indicating that there is a paralogue specific mechanism to adjust the STT3 subunits to the changing needs of the cell during ER stress. This forced degradation of STT3B was partially rescued by RHBDL4 knockdown (Fig. 6B), suggesting that under these conditions also RHBDL4 contributes to tuning of the post-translational glycosylation machinery.

**Fig. 6.**
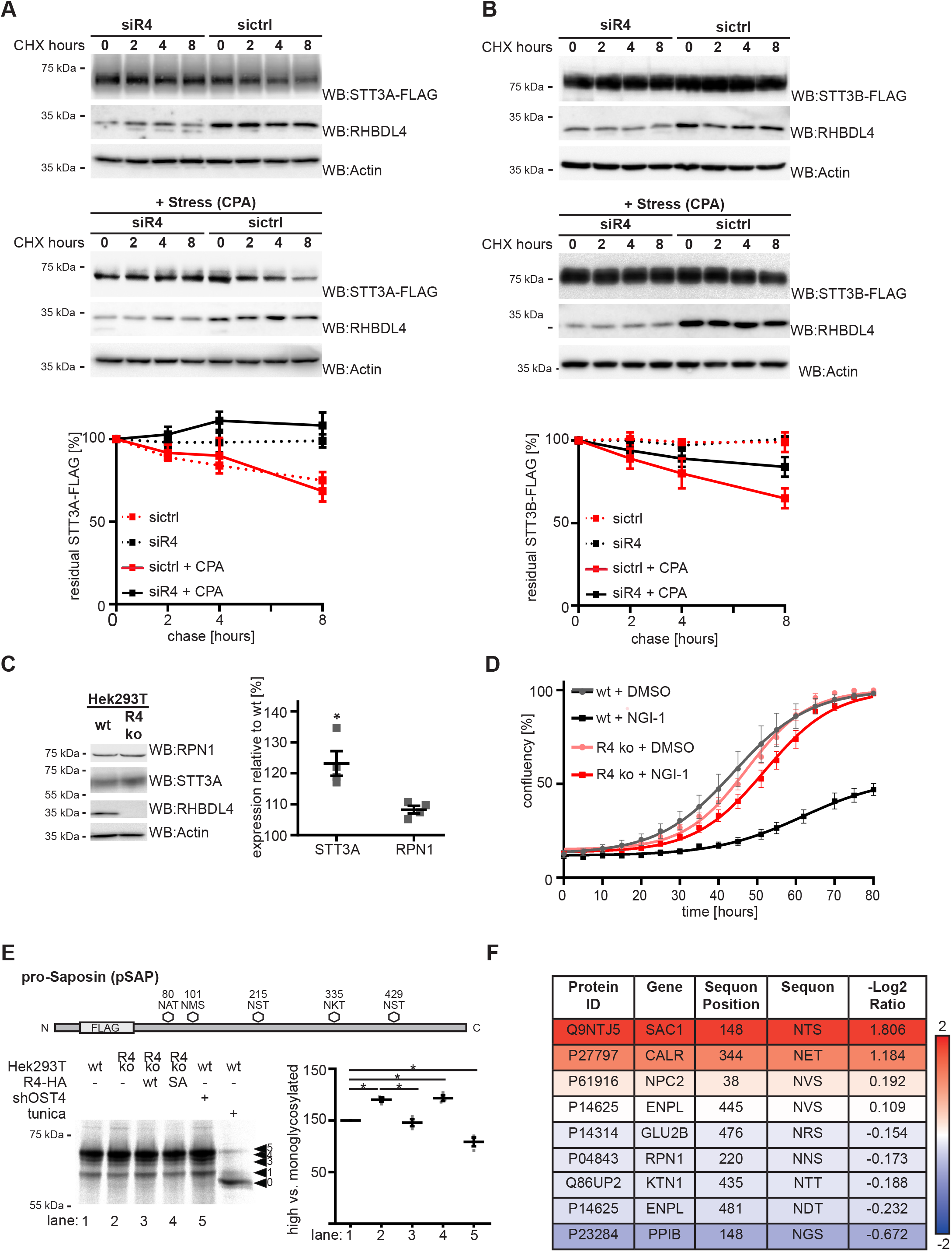
RHBDL4 knockout impacts glycosylation efficiency. (A) STT3A-FLAG expressed from the endogenous locus in Hek293T cells is stabilized after depletion of RHBDL4 when compared to with control, as shown by cycloheximide (CHX) chase. Degradation can be further enhanced by treatment with cyclopiazonic acid (CPA, 1 μM) 24 hours prior to addition of CHX. Actin was used as loading control. Actin levels are shown as loading control. Below quantification of STT3A-FLAG expression level relative to respective timepoint 0 (means ± SEM, n=3). (B) STT3B-FLAG expressed from the endogenous locus in Hek293T cells is stable within 8 hours as shown by CHX chase and degradation is induced by treatment with CPA 24 hours before the chase experiment. Actin levels are shown as loading control. Below quantification of STT3B-FLAG expression level relative to respective timepoint 0 (means ± SEM, n=3). (C) Western blot (WB) analysis of endogenous STT3A and RPN1 in total cell lysates from Hek293T wt or RHBDL4 knock out cells (R4 ko) cells. Actin levels are shown as loading control. Right panel, western blot quantification (means ± SEM, n=4, *p < 0.05, student’s T-test). (D) Growth curves of Hek293T wt and R4 ko cells treated with OST inhibitor NGI-1 (10 μM) or vehicle control (DMSO). Mean confluency of three technical replicates is shown (± SEM). Line represents sigmoidal fitted curve for the respective condition. (E) Hek293T wt and R4ko cells were transfected with glycosylation reporter prosaposin (pSAP-FLAG) and fraction of FLAG-tagged pSAP glycosylated at multiple sites was determined by 10 min radioactive pulse-labelling, immunoprecipitation and autoradiography. Right panel shows quantification of highly glycosylated pSAP relative to monoglycosylated form (means ± SEM, n=5). Ectopic expression of HA-tagged RHBDL4 (HA-R4) rescues reduction observed in R4ko cells whereas expressing either empty vector (−) or catalytically inactive SA mutant did not. In contrast, shRNA targeting of OST4 that destabilizes STT3A increases the mono-glycosylated pSAP species. Pretreatment of cells for 16 hours with tunicamycin completely blocking N-linked glycosylation was used as control. (means ± SEM, n=3; *p < 0.05, one-way ANOVA) (F) Overview of proteins which have been identified to occur as partially glycosylated and with glycan occupancy ratio change of greater than 0.1 between HEK292T wt and R4ko cells. Positive values indicate increased glycosylation compared to wt. Refer to methods section for detailed description of the applied analyses steps.

We next asked whether the RHBDL4-mediated degradation of OST subunits in untreated cells has an effect on their steady state levels. In other words, can RHBDL4 alter the abundance of functional OST complexes, thereby modulating the glycosylation capacity of the cell? We generated RHBDL4 knock-out Hek293T cells and confirmed functionality of the knock-out by showing delayed degradation of ectopically expressed RPN1 construct in a cycloheximide chase experiment (Fig. S5A). Next, we analyzed steady state levels of endogenous STT3A and RPN1 in our RHBDL4 knock-out cell line. While transcription of both genes is not affected (Fig. S5B), western blot analysis of the endogenous proteins showed that steady-state levels are increased upon loss of RHBDL4, although for RPN1 this failed to reach significance (Fig. 6C). This increase in available active OST subunits results in a fitness benefit of RHBDL4 knock-out Hek293T cells compared to wt cells after treatment with the N-glycosylation inhibitor NGI-1, a dose leading to partial reduction of N-linked glycosylation (Fig. 6D and S5C and D) (Lopez-Sambrooks et al., 2016), as assessed by monitoring the glycosylation state of SPP (Fig. S5C) (Weihofen et al., 2000). That increase in available active OST subunits indeed leads to an increased glycosylation capacity was demonstrated by monitoring glycosylation of the STT3A reporter protein proSaposin (pSAP) (Ruiz-Canada et al., 2009). In control cells, the predominant form of ectopically expressed pSAP is the fully-glycosylated form, while tunicamycin treatment effectively prevented any N-glycosylation and yielded non-glycosylated pSAP (Fig. 6E). As previously observed, depletion of OST4 (Dumax-Vorzet et al., 2013) resulted in a substantial perturbation of pSAP N-glycosylation with an increase of the monoglycosylated form. Loss of RHBDL4 led to a decrease of monoglycosylated relative to multi-glycosylated pSAP that could be rescued by transfection of RHBDL4 wt but not SA. Although these effects are small, they are consistent with the ~20% increase in STT3A steady state level upon knock-out of RHBDL4 (Fig. 6C). Consistent with this subtle increase of OST levels, MS-analysis of the N-gycoproteome of RHBDL4 knockout cells revealed increased glycosylation of the partially glycosylated ER proteins. To determine site-specific *N*-glycan occupancy we used a ratiometirc approach (Xu et al., 2015; Zhu et al., 2014) where we compared overall protein abundance to the abundance of non-glycosylated sequon-containing peptides. As treatment with PNGase F, which cleaves between the innermost GlcNAc and asparagine residues of *N*-linked glycans, converts previously glycosylated asparagines to aspartic acids the resulting mass shift can be used to assign peptides as unmodified or occupied. We identified 219 peptides with deamidation events, suggesting they were previously glycan occupied. The majority of sequons identified were already fully glycosylated in Hek293T wt cells or did not show a log2 change in glycosylation occupancy of greater than 0.1 (Table S3-S6). Yet, four ER proteins, namely phosphatidylinositide phosphatase SAC1, calreticulin (CALR), NPC intracellular cholesterol transporter 2 (NPC2) and endoplasmin (ENPL) (Fig. 6F) showed a decrease of the abundance of unglycosylated peptides upon loss of RHBDL4, suggesting increased glycosylation in the RHBDL4 knock-out cell compared to wt. Taken together, these results underline the impact of increased OST levels in our tissue culture model. The molecular mechanisms leading to a basal and stress-induced turnover, and the physiological function in tuning the ER glycosylation capacity by RHBDL4, are important questions that remain to be addressed.

## Discussion

With the aim of characterizing the physiological function of RHBDL4, we used substrate trapping in combination with CRISPR-based chromosomal tagging to identify and validate endogenous RHBDL4 substrates. With the discovery of several OST subunits including STT3A and STT3B as native RHBDL4 substrates, we confirm our previous finding relying on mutated ERAD substrates, that polytopic membrane proteins are targeted for rhomboid-dependent ERAD (Fleig et al., 2012) and extend it to endogenously occurring substrates. Different to rhomboids in the Golgi and late secretory pathway, which primarily act as protein secretases (Lichtenthaler et al., 2018), we observe that RHBDL4-catalyzed cleavage triggers degradation of OST subunits. This function in protein catabolism goes along with a mechanism that allows RHBDL4 to select specifically a designated fraction of the entire substrate population for cleavage. Likewise, we showed that this ERAD substrate clipping mechanism acts on OST subunits when they are orphan; either expressed in excess or due to reduced expression of other complex subunits. In addition, chemically-induced ER stress can trigger RHBDL4-mediated degradation of STT3B. Therefore, RHBDL4 influences OST complex homeostasis in a subunit-specific and potentially regulatable manner. Effects on the endogenous substrates that we observe upon RHBDL4 inhibition seem to be moderate only. Yet, in the light of degradation as a regulatory mechanism such subtle abundance changes are predicted to have primarily a modulatory effect acting faster than other homeostasis mechanisms such as transcription. Importantly, loss of RHBDL4 leads to increased stability of STT3A, resulting in an overall increased glycosylation capacity of Hek293T cells. Increased tolerance of RHBDL4 knockout cells to the STT3 inhibitor NGI-1 (Lopez-Sambrooks et al., 2016) demonstrates that even subtle OST abundance changes can have a profound impact on cellular physiology.

Previously, we demonstrated a role for RHBLD4 in ER protein homeostasis by facilitating ERAD of membrane proteins with unstable TM domains (Fleig et al., 2012). More recently, it has been shown that RHBDL4 can process Alzheimer’s disease-associated APP (Paschkowsky et al., 2016). We now show that the physiological substrate spectrum of RHBDL4 also includes native polytypic membrane proteins such as STT3A and STT3B. While under steady-state conditions the majority of these OST components are stable, upon ectopic overexpression their assembly into protein complexes is saturated and orphan subunits are created. Unpaired OST complex partners might be deregulated in their function or cause harm to other proteins by exposing otherwise masked hydrophobic patches in the lumen or charges in the TM region (Juszkiewicz and Hegde, 2018). Severe protein complex stoichiometry imbalance is observed in aneuploid cells leading to induction of stress response pathways (Brennan et al., 2019; Oromendia et al., 2012). Since aneuploidy is a hallmark of cancer cells, the need for RHBDL4-mediated removal of surplus complex subunits may explain why the *RHBDL4/Rhbdd1* gene is transcriptional upregulated in various forms of tumors (Song et al., 2015; Zhao et al., 2019). RHBDL4 efficiently cleaves these orphans thereby accelerating their disposal by the ERAD dislocation machinery. This is consistent with earlier work in yeast showing that ERAD compensates for disturbed OST complex homeostasis by degrading excess subunits (Mueller et al., 2015). While in yeast this is mediated by the Asi E3 ligase complex located in the inner nuclear membrane (Natarajan et al., 2019), mammalian cells do not have Asi homologues and at least partially rely on RHBDL4 mediated ERAD.

Furthermore, we and others previously showed that positive charged TM residues trigger recognition of ERAD-M substrates by RHBDL4 (Fleig et al., 2012; Recinto et al., 2018). Consistent with this, we identify several positive charges in STT3A positioned on the interface of the catalytic subunit and the lipid carbonyl region that may therefore be recognized by RHBDL4. In general, ERAD-M degrons appear to be characterized by metastable TM segments that are subject to several different quality control mechanisms including but not limited to recognition by the Hrd1 and Asi2 ERAD E3 ubiquitin ligases (Natarajan et al., 2019; Sato et al., 2009), intramembrane proteolysis by SPP (Yucel et al., 2019), an assembly-dependent membrane integration of nascent TM segments at the Sec61 channel and the ER membrane protein complex (Chitwood et al., 2018; Coelho et al., 2019; Feige and Hendershot, 2013; Tector and Hartl, 1999). Additionally, rhomboid family proteins, with their ability to specifically recognize TM segments, emerge as important safeguards of the membrane proteome (Lemberg, 2013). In this regard the best characterized is the bacterial GlpG, which recognizes a combination of TM helix dynamics and primary amino acids sequence surrounding the cleavage site (Akiyama and Maegawa, 2007; Strisovsky et al., 2009). Substrate TM segments are thought to first bind to a putative membrane-embedded docking site from where they unfold into the active site (Strisovsky, 2016). We show here that a conserved negative charged cluster in the active site gating TM segment H2 consisting of two conserved aspartate residues (D77/D78, see Fig. S4E and F) is important for efficient cleavage of STT3A but not of STT3B. Based on this result, we suggest that the positive charged TM residues in STT3A may be recognized by such a putative docking site that is at least partially formed by this feature. It remains to be shown whether this recognition involves unfolding of substrate TM helices or whether the substrate TM helix remains bound to the rhomboid protease exosite while luminal loops exposing cleavage site regions enter the active site groove. Mapping of the cleavage sites revealed that the scissile peptide bonds in L1 and L9 are spaced apart from the TM helix charge, supporting a model that docking and formation of the scission complex are distinct steps. Overall, the emerging picture is that recognition of RHBDL4 substrates is influenced by multiple factors, also including substrate ubiquitination by ERAD E3 ligases (Fleig et al., 2012), integration of signaling from G-protein coupled receptors (Wunderle et al., 2016), tyrosine phosphorylation of the RHBDL4 cytoplasmic domain (Ikeda and Freeman, 2019) and sensing of the membrane lipid composition (Paschkowsky et al., 2018).

As mentioned earlier, the canonical ERAD dislocation pathway also plays an important regulatory role in degradation of functional proteins in response to changing demands of the cell (Christianson and Ye, 2014; Johnson and DeBose-Boyd, 2018; Mehrtash and Hochstrasser, 2018; Ruggiano et al., 2014). However, to date no abundance control mechanism has been described that specifically disassembles multimeric TM protein complexes in the ER. Here, we provide evidence that RHBDL4 induces degradation of the two OST catalytic subunits STT3A and STT3B, even when expressed at endogenous levels, indicating that they might also be recognized while native complex assembly occurs. For STT3A this degradation occurs in the absence of any stimulus whereas OST complexes harboring STT3B seem to be more stable and are only degraded upon ER stress. Furthermore, within STT3A, both cleavage sites (L1 and L9) are surface exposed (Fig. 3B) (Braunger et al., 2018), indicating that RHBDL4 could cleave its substrates even in the context of mature complexes. It remains an important question whether this is a mechanism that allows to inactivate functional OST complexes. Why are these proteins cleaved by RHBDL4 prior to recognition by the ERAD dislocation machinery and turnover by the proteasome? Cleavage in L1 and L9 splits the STT3 active site into sperate polypeptide chains and for the non-catalytic OST subunits RHBDL4-catalyzed clipping disintegrates important functional properties from the TM anchor. We may speculate that because in the cellular context proteolysis is an irreversible reaction, RHBDL4 cleavage of OST subunits is more potent in shifting the equilibrium to degradation than the classical dislocation pathway.

A similar role for intramembrane proteolysis in the abundance control of glycosyltransferases has been described previously for the Golgi-resident SPP-like protease 3 (SPPL3) (Voss et al., 2014). This indicates that intramembrane proteolysis plays a global role in the control of protein glycosylation, which has a wide range of important functions such as controlling enzyme activity, protein-protein interactions and protein stability (Cherepanova et al., 2016). In the ER N-linked glycosylation is crucial during protein folding and quality control, as glycans allow entry into the calnexin/calreticulin-mediated folding cycle and eventually target terminally misfolded proteins for ERAD (Ellgaard et al., 2016). Critically however, synthesis of glycans is an energetically expensive process and therefore must be finely tuned to the actual need of the cell. Although there is an initial upregulation of STT3A, STT3B and RPN1 during ER stress (Bergmann et al., 2018), the rate of ER protein synthesis and therefore the need for glycosylation might be reduced over time. As we previously observed that RHBDL4 is upregulated during ER stress (Fleig et al., 2012), we speculate that RHBDL4-catalyzed degradation allows for a fast adaptation of the available OST machinery during the different stages of the UPR. We showed that turnover of STT3A and STT3B, the two alternative OST core components in an unstressed and stressed state is subunit specific. Whereas STT3A is constantly turned over by an RHBDL4-dependent mechanism, STT3B is stable and is only degraded by RHBDL4 upon chemically induced ER stress, meaning the glycosylation activity is shifted towards the co-translationally pathway. An advantage of this putative glycotuning mechanism might be to ensure that only fully glycosylated nascent chains that are subject to calnexin/calreticulin-mediated quality control leave the Sec61 translocon. Alternatively, this altered glycosylation might affect a selected set of proteins, or change OST subunit composition to enhance non-canonical functions, as observed for RPN1 which also acts as a molecular chaperone (Qin et al., 2012). The physiological relevance and molecular mechanism of this subunit specific recognition and degradation by RHBDL4 are important questions that will be part of further investigations.

## Methods

### Plasmids

Unless otherwise stated, all constructs were cloned into pcDNA3.1+ (Invitrogen). Human RHBDL4 (UniGene ID Hs.84236; IMAGE 40023929) was cloned with an N-terminal triple HA-tag. Constructs for C-terminal GFP-tagged human RHBDL4 were generated by subcloning into pEGFP-N1 (Clontech). For generating point mutants, a site-directed mutagenesis strategy was used. For catalytically inactive RHDBL4-SA serine-144 was replaced by alanine and for the RHBDL4-SAUM mutant leucines-274 and leucines-278 where replaced by alanine. Human STT3A (UniGene ID Hs.504237, full-ORF Gateway cDNA clone 111873813), mouse STT3B (UniGene ID Mm.296158, gift from Stephen High) and human B4GAT1 (UniGene ID Hs.8526, full-ORF Gateway cDNA clone 144741302) were fused to an N-terminal triple FLAG-tag. RPN1 (UniGene ID Hs. Hs.518244; IMAGE 3461167), RPN2 (UniGene ID Hs.370895, full-ORF Gateway cDNA clone 107158506), DDOST (UniGene ID Hs.523145, full-ORF Gateway cDNA clone 191432118) and PSAP (UniGene ID Hs.523004, full-ORF Gateway cDNA clone 139868800) were cloned without their signal peptides into a pcDNA3-based expression vector containing the Spitz signal peptide fused to a triple FLAG-tag as has been described (Adrain et al., 2011). STT3A reference peptides were generated by introduction of a premature stop codon at the indicated position. Charge and cleavage site mutants of STT3A where generated by site-directed mutagenesis. For stable expression of GFP-tagged mouse RHBDL4-SA and RHBDL4-SAUIM, the open reading described previously (Fleig et al., 2012) were sub-cloned into pcDNA5/FRT/TO (Invitrogen). For transient knockdown the small hairpin (shRNA)-expressing vector pSUPER.neo targeting OST4 (targeting sequence: 5’-TCAACAATCCCAAGAAGCA-3’) was generated. As non-targeting (nt) control pSUPER.neo targeting 5’-ACAGCUUGAGAGAGCUUUA-3’ designed for knockdown of RHBDL4 in COS7 cells (but not human cells) was used. The sequences of all generated plasmids were verified by DNA sequencing.

### Cell Lines

Hek293T cells (ATCC) were grown in DMEM (Invitrogen) supplemented with 10% fetal bovine serum at 37°C in 5% CO2 and regularly screened for mycoplasma contamination. To prepare inducible stably transfected cells expressing GFP-tagged RHBDL4-SA or RHBDL4-SAUM, pcDNA5/FRT/TO constructs described above were co-transfected with pOG44 (Invitrogen) into Flp-In Hek293 T-REx cells (Invitrogen) followed by selection with blasticidin (10 μg/ml) and hygromycin B (125 μg/ml). Hek293 T-REx cells expressing GFP-tagged mouse RHBDL4 wt were described previously (Fleig et al., 2012). Expression was induced with 1 μg/ml doxycycline 24 hours prior to harvest.

RHBDL4 knockout Hek293T cells were generated using TALEN expression vectors obtained from www.talenlibrary.net (Kim et al., 2013) with the following sequences: left - NGNGNNNGNINGNGNGHDNININGNINGNNNNHDNING-, right - HDNGNGNGHDNGNINNNINGNGNGNINGNGHDHDNGNG-(Kühnle et al., 2019a). Generation of STT3A-FLAG and STT3B-FLAG Hek293T cells with a triple FLAG-tag before the stop codon in the last exon of the respective gene by using CRISPR/Cas12 mediated gene editing has been described before (Fueller et al., 2019). The cells used in this study where generated with the previously described Ctag-STT3A-LbCpf1.rev2 and Ctag-STT3B-LbCpf1.rev2 primer, respectively (Fueller et al., 2019).

### Transfection

Transient transfections were performed using 25 kDa linear polyethylenimine (Polysciences) (Durocher et al., 2002). Typically, 1000 ng (for FLAG-STT3A) to 500 ng (for all other tested substrates e.g. FLAG-RPN1 and FLAG-STT3B) plasmid encoding substrate and 100 ng plasmid encoding RHBDL4 were used per well of a 6-well plate. Total transfected DNA (2 μg/well) was held constant by addition of empty plasmid. For inhibitor studies, MG132 (Calbiochem), diluted in DMSO, was added to the cells 16 hours prior to harvest. Cells were harvested typically 48 hours post transfection. Cells were solubilized in SDS-sample buffer, incubated at 37°C for 30 minutes and analyzed by SDS-PAGE. For RHBDL4 and OST4 depletion 50 pmol (RHBDL4) or 25 pmol (OST4) ON-TARGETplus SMARTpool human siRNA (Dharmacon) and 2.5 μl RNAimax (Invitrogen) transfection reagent per well of a 12-well plate were used. Amount of siRNA was kept constant within an experiment by addition of scrambled control.

### Mass Spectrometry based Substrate Screen

For isolation of endogenous RHBDL4 interaction partners by shotgun proteomics, Hek293 T-REx cells stably expressing mouse GFP tagged RHBDL4 variants (see above) were grown for six doublings in medium containing unlabeled amino acids (RHBDL4 wt, “light”), or supplemented with 13C15N-arginine and 13C15N-lysine (Silantes) (RHBDL4-SA, “heavy”) or 13C-arginine and D4-lysine (RHBDL4-SAUM, “medium”). Expression was induced with 1 μg/ml doxycycline for 24 h. Equal numbers of cells from all three cultures were pooled and microsome fraction was isolated by hypotonic swelling and centrifugation as previously described (Chen et al., 2014). For immunoprecipitation of RHBDL4-GFP, microsomes were solubilized with 1% Triton X-100 in IP buffer, containing complete proteasome inhibitor (Roche) and 10 μg/ml PMSF. Cell lysates were cleared by centrifugation at 10,000 g for 10 min. Pre-clearing and anti-GFP immunoprecipitation was performed as described below. The immunocomplexes were eluted in SDS sample buffer and resolved by SDS-PAGE. Proteins were reduced with dithiothreitol, alkylated with iodacedamide and in gel digested with trypsin. Peptides were extracted with 50% acetonitrile/0.1% trifluoroacetic acid. Samples were analyzed by a UPLC system (nanoAcquity) coupled to an ESI LTQ Oribitrap mass spectrometer (Thermo). The uninterpreted MS/MS spectra were searched against the SwissProt-human database using MaxQuant software. The algorithm was set to use trypsin as enzyme, allowing at maximum for two missed cleavage site and assuming carbamidomethyl as a fixed modification of cysteine, and oxidized methionine and deamidation of asparagine and glutamine as variable modifications. Mass tolerance was set to 4.5 ppm and 0.2 Da for MS and MS/MS, respectively. In MaxQuant the “requantify” and “match between runs” option was utilized, the target decoy method was used to determine 1% false discovery rate. For generation of the substrate candidate list the following filters where applied (compare Table S1 and S2): quantified in both replicates, more or equal to 2 unique peptides per replicate, gene ontology cellular compartment annotation contains “integral to membrane” and “endoplasmic reticulum” and the mean heavy over light ratio is greater or equal 2.0.

### Glycoproteomics

For detection of N-linked glycosylation sites, approximately 20 μg of Hek293T R4ko cells was denatured and reduced by incubation in 6 M guanidine-HCl, 50 mM TrisHCl buffer pH 8, and 10 mM dithiothreitol (n=2). Reduced proteins were alkylated with 25 mM acrylamide and the reaction was quenched with 5 mM dithiothreitol followed by precipitation at −20°C in four volumes of methanol/acetone (1:1 v/v). The precipitated proteins were centrifuged for 10 min at 18,000 g and the protein pellet resuspended 50 mM NH_4_HCO_3_ before digestion with sequencing grade porcine trypsin (enzyme to protein ratio of 1:40) (Sigma-Aldrich, MO, USA) for 16 hours at 37°C. From each sample approximately 10 μg of protein was taken and pooled, then deglycosylated with 2000 U of Peptide-N-Glycosidase F (PNGase F) (New England BioLabs, MA, USA). All samples were incubated for 16 hours at 37°C. The pooled deglycosylated samples were then subjected to high pH fractionation (Enculescu et al., 2019) and eluted into 8 fractions before drying and reconstitution in 0.1% formic acid. The tryptic peptides from each replicate (not treated with PNGase F) were desalted and concentrated with C18 ZipTips (10 μl pipette tip with a 0.6 μl resin bed; Millipore, MA, USA), before drying and reconstitution 100 μl 0.1% formic acid. Samples were analyzed by LC-ESI-MS/MS using a Shimadzu Prominence nanoLC system coupled to a TripleTof 5600 instrument (ABSciex) using a Nanospray III interface as previously described (Zacchi and Schulz, 2016). For data-dependent acquisition (DDA) 30 μl of each high pH fraction was injected. For sequential window acquisition of all theoretical mass spectra (SWATH) analysis 8 μl (~400 ng) of the tryptic peptides not treated with PNGase F were injected. Data produced from the DDA analyses were searched in ProteinPilot v5.0.1 using a protein database containing the UniProt reference human proteome (Proteome ID UP000005640, download on 20. April 2018) and a custom contaminants database. Standard settings included: sample type, identification; digestion, trypsin; instrument, TripleTof 5600; cysteine alkylation, acrylamide; search effort, thorough. False discovery rate analysis using ProteinPilot was performed on all searches. Peptides identified with greater than 99% confidence and with a local false discovery rate of less than 1% were included for further analyses. From the ProteinPilot search results (Table S3-S5), distinct peptides were searched using N-glycosylation Sequon Parser (http://schulzlab.glycoproteo.me/#/nsp) to identify peptides containing N-linked sequons (asparagine-Xaa-serine/threonine where Xaa is not proline). The ProteinPilot data was also used as an ion library to measure peptide abundance using PeakVeiw v2.1 (SCIEX) with the settings: shared peptides, allowed; peptide confidence threshold, 99%; false discovery rate, 1%; exclude modified peptides, enabled; XIC window, 6 min; XIC width, 75 ppm. The results were searched for peptides containing N-linked sequons and the amino acid position of the potential N-linked site in the protein sequence was annotated. In addition, for each protein identified in the results the subcellular location was retrieved from UniProt. As *N*-glycosylation events within the samples were confirmed by treating tryptic peptides with PNGase F, which cleaves between the innermost GlcNAc and asparagine residues of *N*-linked glycans, previously glycosylated asparagine was converted to aspartic acid. Accordingly, asparagin residues within *N*-linked consensus sites can be assigned as unmodified or occupied. This relative peptide abundances were used to determine site-specific N-linked occupancy by calculating the ratios of the intensities of each unmodified (not glycosylated) sequon containing peptide to the corresponding protein (Table S6) (Xu et al., 2015). The mean ratios of RHBDL4 knock-out was divided by RHBDL4 wt corresponding ratios and the - log2 ratio calculated. Consequently, negative ratios depict a relative increase of glycan site occupancy in knock-out versus wt condition. Results where filtered for log2 changes of equal or greater than 0.1.

### Pulse-label Analysis

For pulse-label experiments, transfected Hek293T cells were starved for 60 min in methionine/cysteine free DMEM (Invitrogen) supplemented with 10% dialyzed fetal calf serum and then metabolically labelled for 10 min with 55 μCi/ml ^35^S-methionine/cysteine protein labeling mix (Perkin Elmer). Cells were rinsed immediately with PBS twice and solubilized with 1% Triton X-100 in solubilization buffer (50 mM HEPES-KOH, pH 7.4, 150 mM NaCl, 2 mM MgOAc_2_, 10% glycerol, 1 mM EGTA), containing EDTA-free complete protease inhibitor cocktail (Roche) and 10 μg/ml PMSF followed by immunoprecipitation of FLAG-tagged proteins as described below. Samples were resolved by SDS-PAGE and labeled proteins were visualized by a FLA-7000 phosphorimager (Fuji). Image quantification was performed using Fiji (Rasband, W.S., ImageJ, U. S. National Institutes of Health, Bethesda, Maryland, USA, http://imagej.nih.gov/ij/, 1997-2014) (Schindelin et al., 2012).

### Immunoprecipitation

For immunoprecipitation of GFP tagged proteins, cells were solubilized with solubilization buffer (50 mM HEPES-KOH, pH 7.4, 150 mM NaCl, 2 mM MgOAc_2_, 10% glycerol, 1 mM EGTA), containing 1% Triton X-100, EDTA-free complete protease inhibitor cocktail (Roche) and 10 μg/ml PMSF. Cell lysates were cleared by centrifugation at 15,000 g for 15 min and subsequently pre-incubated for 1 h on BSA-coupled sepharose beads. Anti-GFP immunoprecipitation was performed using a GFP-specific single chain antibody fragment coupled to NHS-activated sepharose beads as described before (Fleig et al., 2012). Immunoprecipitates were washed three times in solubilization buffer, containing 0.1% Triton X-100 and eluted in SDS sample buffer and analyzed by western blotting (see below).

For immunoprecipitation of FLAG tagged proteins lysates were prepared as described above. Anti-FLAG immunoprecipitation was performed using anti-FLAG agarose beads (M2, Sigma). Immunoprecipitates were washed two times in solubilization buffer containing 0.1% Triton X-100 and eluted in SDS sample buffer and analyzed by western blotting (see below).

### Cycloheximide chase

Cycloheximide (100 μg/ml) chase was conducted 24 h after transfection of Hek293T cells and cell extracts were subjected to western blot analysis as described below.

### Antibodies

The following antibodies were used in the mentioned dilutions: 1:1000 mouse monoclonal anti-β-actin (Sigma, A1978), 1:1000 mouse monoclonal anti-FLAG (M2; Sigma, F3165), 1:1000 mouse monoclonal anti-FLAG-HRP (M2; Sigma, A8592), 1:1000 mouse monoclonal anti-GFP (Roche, 11814460001), 1:1000 mouse monoclonal anti-HA (3F10; Roche, 11867423001), 1:1000 rabbit polyclonal anti-RPN1 (abcam, ab198508), 1:500 rabbit polyclonal anti-STT3A (Proteintech, 12034-1-AP), 1:1000 rabbit polyclonal anti-SPP (gift from Chica Schaller), 1:250 rabbit polyclonal anti-RHBDL4 (Sigma, HPA013972) and 1:1000 as previously described (Fleig et al., 2012).

### Western Blotting

Total cells, lysates or immunoprecipitates were solubilized in SDS sample buffer (50 mM Tris-Cl pH 6.8; 10 mM EDTA, 5% glycerol, 2% SDS, 0.01% bromphenol blue) containing 5% β-mercaptoethanol. For Endo H digest prior to SDS-PAGE, samples in SDS containing sample buffer have been diluted with water to contain 0.5 % SDS and have been digested with Endo H (NEB) according to instructions by the manufacturer. For western blot analysis samples were incubated for 30 min at 37°C, resolved on Tris-glycine SDS-PAGE, transferred on PVDF membrane, followed by western blot analysis using enhanced chemiluminescence to detect bound antibodies. For detection the LAS-4000 system (Fuji) was used and Fiji (Schindelin et al., 2012) was used for image quantification. Data shown are representative of three independent experiments.

### Homology Model

The homology model of the membrane integral part of RHBDL4 was built using the Phyre2 web portal for protein modeling, prediction and analysis (Kelley et al., 2015). The following residues of the RHBDL4 sequence (UniProt Q8TEB9) were included in the homology model: 25-123; 137-211 in order to restrict the modeling to all sequence parts which showed a homology confidence of 100%.

### Real time quantitative reverse transcription and polymerase chain reaction

RNA was isolated using NucleoSpin RNA kit (Machery-Nagel) according to the manufacturer’s description. Complementary DNA (cDNA) transcripts were generated using the RevertAid First Strand cDNA Synthesis Kit (Thermo Fisher) by use of the accompanied random hexamer primer. Amplification reactions where performed using the SensiFAST SYBR No-ROX Kit (Bioline) according to the manufacturer’s instructions. β-2 microglobulin (B2M) and TATA-binding protein (TBP) have been used for normalization. The following primers have been used: B2M: CACGTCATCCAGCAGAGAAT, TGCTGCTTACATGTCTCGAT; TBP: CCGGCTGTTTAACTTCGCTT, ACGCCAAGAAACAGTGATGC; OST4: CGTTCTCTATCACTACGTGGC, GTGATGGGTAGAAGGGATGAAG; RPN1: GACATTGTGGTCCACTACACG, AACAGGATGTAGAAGGCCGC; STT3A: CCAGGCATTCCAGAAAACAGC, AACACACGACGATCAGTTTGC; STT3B: AAAGAAGACTACCAAAAGGAAGCG, CAAGTTAGGAACCAGAACAGTGC. All reactions were performed in technical triplicate using 384-well plates on the LightCycler 480 System using the thermal cycling conditions as per the manufacturer’s instructions (Roche Diagnostics GmbH). The 2^DDCt method was used to calculate relative changes in gene expression and normalization to the arithmetic mean of both B2M and TBP.

### Kinetic growth assay

Cells were harvested by trypsinization, counted and plated at 4000 cells per well in 96-well tissue culture plates in 3 technical replicates. 16 hours after seeding, a sub-effective concentration of NGI-1 (Sigma) (10 μM, IC_50_ approx. 1 μM) from a 10 mM stock in DMSO was added and acquisition of photomicrographs using an Incucyte live cell imager (Essen Biosciences) was started. Images with a 10-fold magnification were acquired over the course of 80 hours. Confluence of the cultures was measured using Incucyte software (Essen Biosciences). Out of focus images were manually removed from the analysis. One representative result of three independent biological replicates is shown.

### Quantification and statistical analysis

Assays were conducted at least in triplicate, with standard error of means (SEM) reported. Quantification of steady-state and cycloheximide chase experiment is compared to the respective control condition or time point zero. For quantification of cleavage efficiency, amount of cleavage product is compared cleavage product in control condition. Statistical analysis was performed using GraphPad Prism version 6 for Windows (GraphPad Software).

## Abbreviations

ERAD: ER-associated degradation
OST: oligosacharyltransferase
TM: transmembrane
UIM: ubiquitin-interacting motif

## Acknowledgments

We thank Stephen High for reagents, Sabine Strahl, Colin Adrain and Stefan Pfeffer for critical reading of the manuscript and Thomas Ruppert (ZMBH MS facility) for MS/MS analysis. The work was supported by funds from the Deutsche Forschungsgemeinschaft (SFB 1036, TP 12) to MKL and a fellowship of the Boehringer Ingelheim Fonds to JDK.

## Author Contributions

J.D.K. designed and performed most experiments and wrote the manuscript. N.L. performed the SILAC substrate proteomics screen. C.P and B.S. performed the glycoproteomics and analyzed data. N.K. generated and validated the RHBDL4 knockout cells, C.W.W. and S.H. helped validating substrate candidates and M.K.L. guided the project, designed experiments, and wrote the manuscript.

