## supplementary material for "Intramembrane protease RHBDL4 cleaves oligosaccharyltransferase subunits to target them for ER-associated degradation"

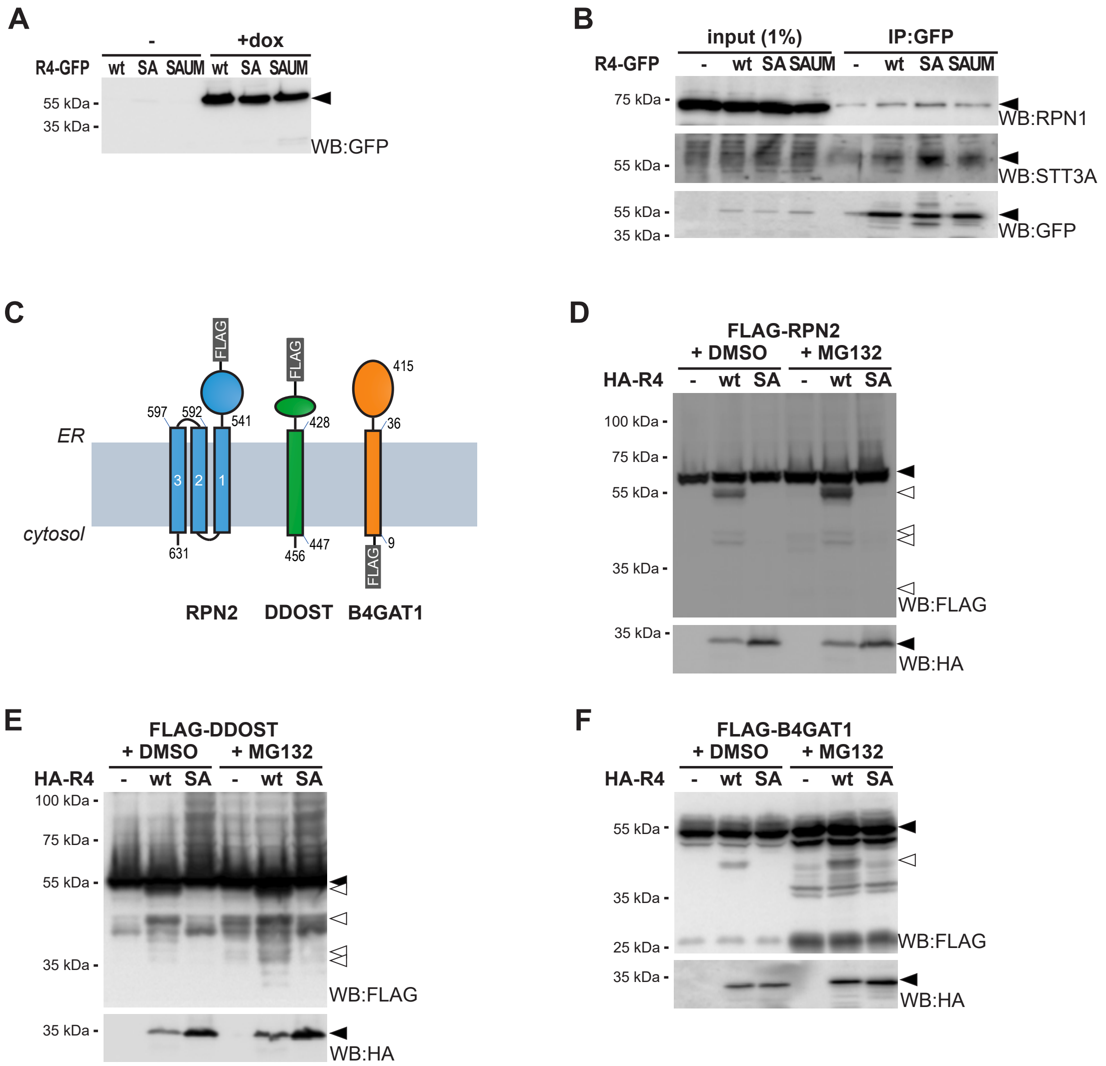


**Fig. S1. Validation of MS Screen and additional RHBDL4 substrate candidates.** (A) Hek293 T-REx cells stably express C-terminally GFP tagged RHBDL4 (R4-GFP) wt, catalytically inactive SA or catalytically inactive RHBDL4 with an additional mutated UIM (SAUM) upon induction with doxycycline (dox). WB, western blot. (B) Hek293T cells transiently transfected with empty vector (-), R4-GFP wt, the catalytically inactive SA mutant or the SAUM double mutant were lysed with Triton X-100 and subjected to GFP-specific immunoprecipitation (IP). Western blots (WB) analysis reveals increased co-purification of endogenous RPN1 and STT3A. (C) Outline of N-terminally FLAG-tagged FLAG-RPN2, FLAG-DDOST and FLAG-B4GAT1 constructs used in this study. Numbers indicate position of TM domain boundaries. (D) Hek293T cells were co-transfected with FLAG-RPN2 and either an empty vector (-), N-terminally HA-tagged RHBDL4 (HA-R4) wt or the catalytically inactive SA mutant. RHBDL4 generates N-terminal cleavage fragments (open triangle) that are degraded by the proteasome as shown by increased steady state level upon MG132 treatment (2 µM) compared to vehicle control (DMSO). (E) Cleavage assay as shown in (D) but with FLAG-DDOST as substrate. (F) Cleavage assay as shown in (D) but with FLAG-B4GAT1 as substrate.

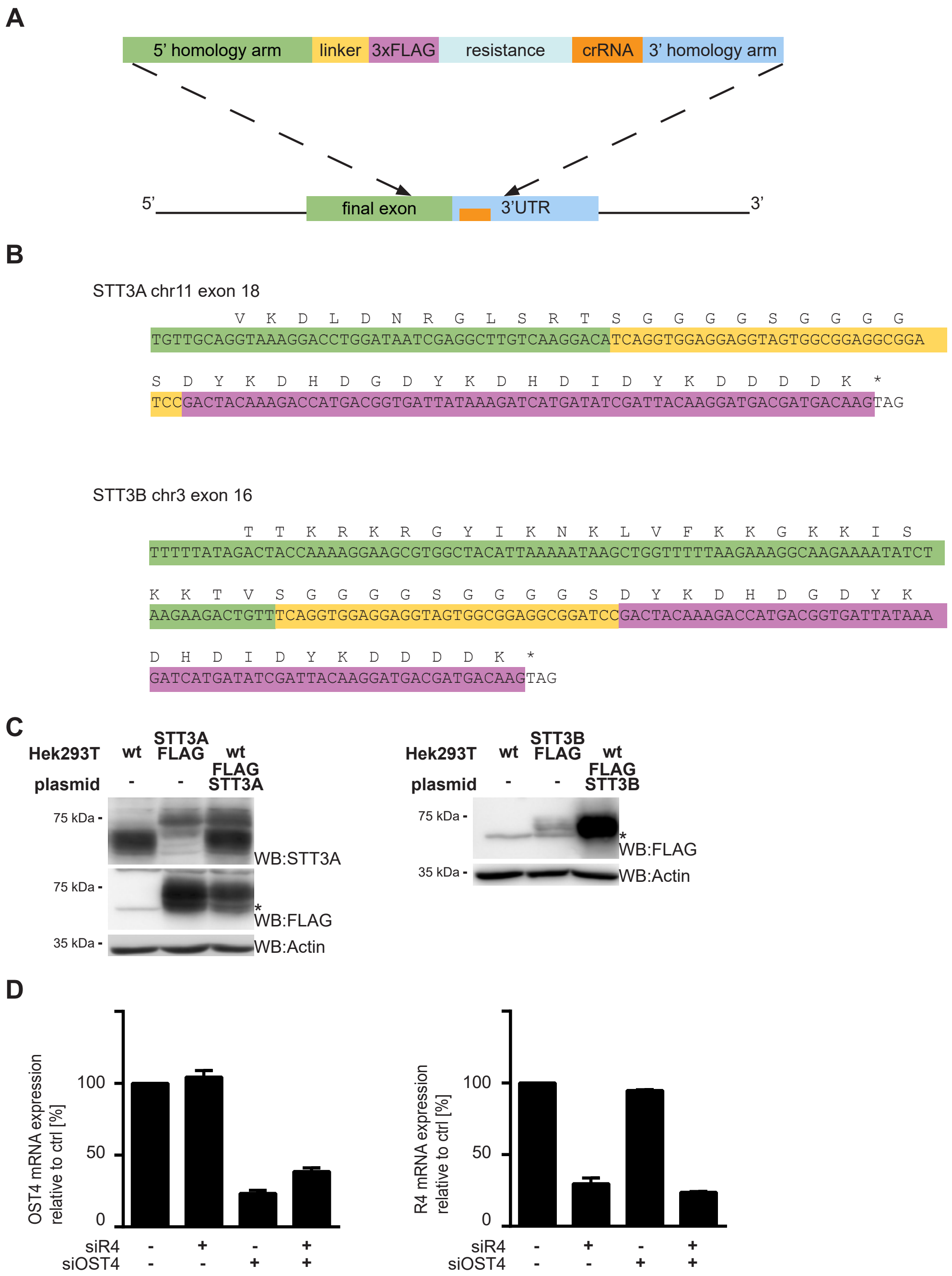


**Fig. S2. Endogenously tagged STT3A and STT3B.** (A) Outline of the applied tagging strategy according to (Fueller et al., 2019). (B) Sanger sequencing of chromosomal DNA obtained from the indicated Hek293T cell lines shows insertion of the triple-FLAG tag in the last coding exon. Color code as in (A). (C) Western blot (WB) analysis of endogenous STT3A, chromosomally tagged STT3A-FLAG and ectopically expressed FLAG-STT3A (left) and WB analysis of chromosomally tagged STT3B-FLAG and ectopically expressed FLAG-STT3B (right) from total cell lysates from Hek293T wt or HEK293T cell line with endogenously tagged STT3A or STT3B, respectively. Empty Vector transfected (-). Ectopic expression of FLAG-STT3A (from 1 µg plasmid) leads to steady state level comparable to the endogenously tagged STT3A-FLAG. 0.5 µg plasmid encoding FLAG-STT3B leads to overexpression as judged by the FLAG-signal. Of note, no STT3B-specific antibody was available restricting the analysis to FLAG-tag. Actin levels are shown as loading control. Asterisk, unspecific band. (D) Transcriptional levels of OST4 and RHBDL4 in HEK293T wt cells of an experiment as shown in Fig. 2A and B relative to sictrl. Normalized to the arithmetic mean of the housekeeping genes β-2 microglobulin and TATA-binding protein.

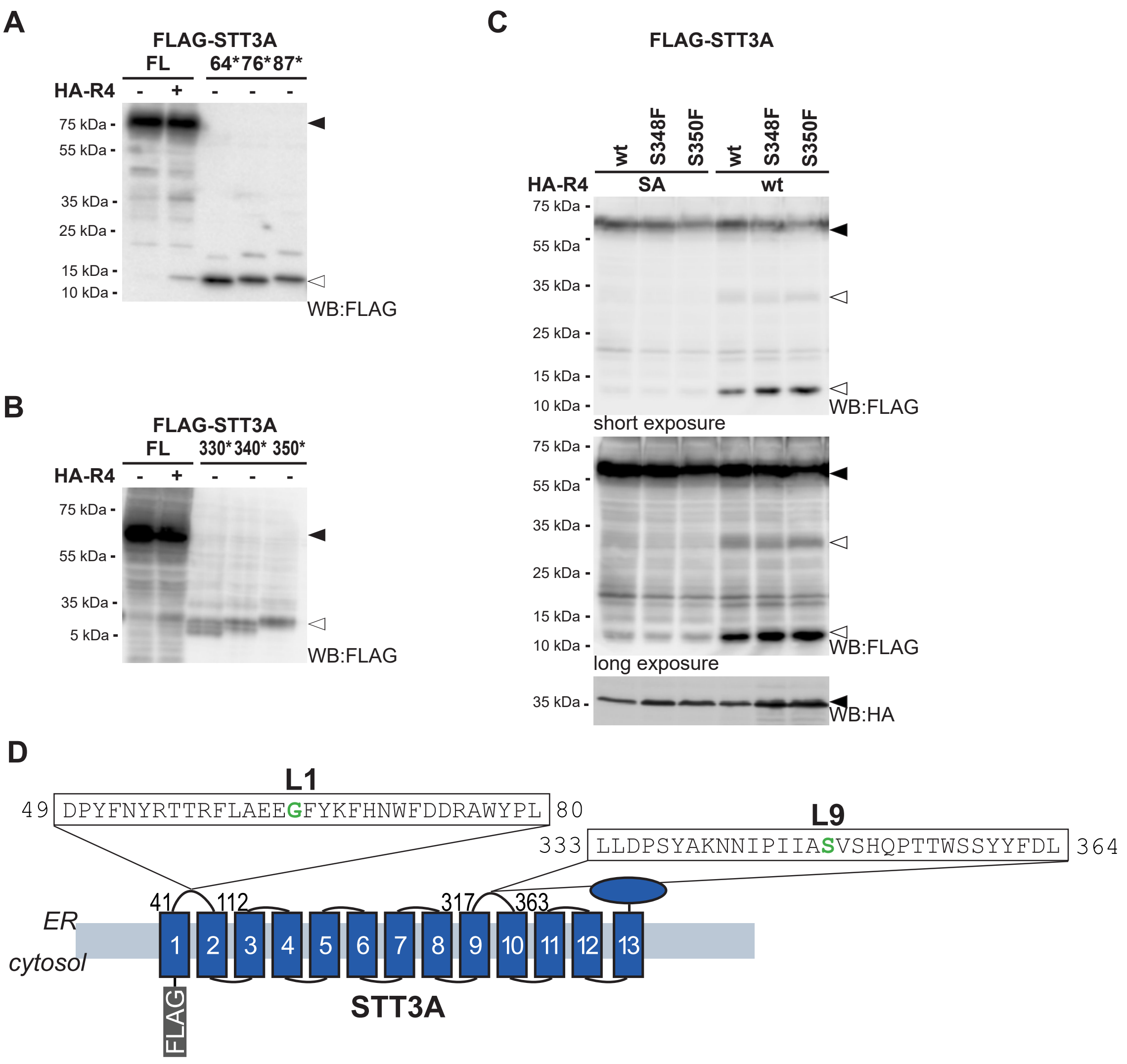


**Fig. S3. STT3A is cleaved within two distinct luminal loops.** (A) Hek293T cells were transfected with full-length (FL) FLAG-STT3A and either an empty vector (-) or HA-tagged RHBDL4 (HA-R4). In addition, cells were transfected with FLAG-STT3A variants truncated at position 64 (64*), 76 (76*) and 87 (87*). Open triangle highlights RHBDL4 generated cleavage fragment at 13 kDa. Cells have been treated with MG132 (2 µM). WB, western blot. (B) Assay as in (A) but FLAG-STT3 with truncations after amino acid 330, 340 and 350 have been used as reference peptides. (C) Hek293T cells were transfected with FLAG-STT3A wt, or with S348F, or a S350F mutant of FLAG-STT3A (filled triangle) together with either the catalytically inactive SA mutant or HA-tagged RHBDL4 (HA-R4) constructs as indicated. Overexpression of the active rhomboid (wt) resulted in cleavage fragments at 35 kDa (L9) and 13 kDa (L1) (open triangles) observed significantly less for the SA mutant. In order to prevent clearance of cleavage fragments, cells have been treated with proteasome inhibitor MG132 (2 µM). (D) Outline of luminal STT3A sequences flanking the predicted RHBDL4 cleavage sites at G64 and S348. Numbers indicate position of TM domain boundaries.

**A**

```
TMD1  STT3A LLKLLILSMAAVLSFSTRLFAVLRF
      STT3B LLSFTILFLAWLAGFSSRLFAVIRF

TMD2  STT3A IRNVCVFLAPLFSSFTTIVTYHLTK
      STT3B IRDVCVFLAPTFSGLTSISTFLLTR

TMD3  STT3A   DAGAGLLAAAMIAVVPGYISR
      STT3B   NQGAGLLAACFIAIVPGYISR

TMD4  STT3A     IAIFCMLLTYYMWIKAVK
      STT3B     IAIFALQFTYYLWVKSVK

TMD5  STT3A    SICWAAKCALAYFYMVSSW
      STT3B    SVFWTMCCCLSYFYMVSAW

TMD6  STT3A    GYVFLINLIPLHVLVLMLT
      STT3B    GYVFIINLIPLHVFVLLLM

TMD7  STT3A   SHRIYVAYCTVYCLGTILSMQ
      STT3B   SKRVYIAYSTFYIVGLILSMQ

TMD8  STT3A  SSEHMAAFGVFGLCQIHAFVDYL
      STT3B  TSEHMAAAGVFALLQAYAFLQYL

TMD9  STT3A QFEVLFRSVISLVGFVLLTVGALL
      STT3B EFQTLFFLGVSLAAGAVFLSVIYL

TMD10 STT3A     LQLLVFMFPVGLYYCFSN
      STT3B     LHILVCTFPAGLWFCIKN

TMD11 STT3A      ARIFIIMYGVTSMYFS
      STT3B      ERVFVALYAISAVYFA

TMD12 STT3A VRLMLVLAPVMCILSGIGVSQVLSTYM
      STT3B VRLMLTLTPVVCMLSAIAFSNVFEHYL

TMD13 STT3A  VASGMILVMAFFLITYTFHSTWV
      STT3B  IKSIVTMLMLMLLMMFAVHCTWV
```

**B**

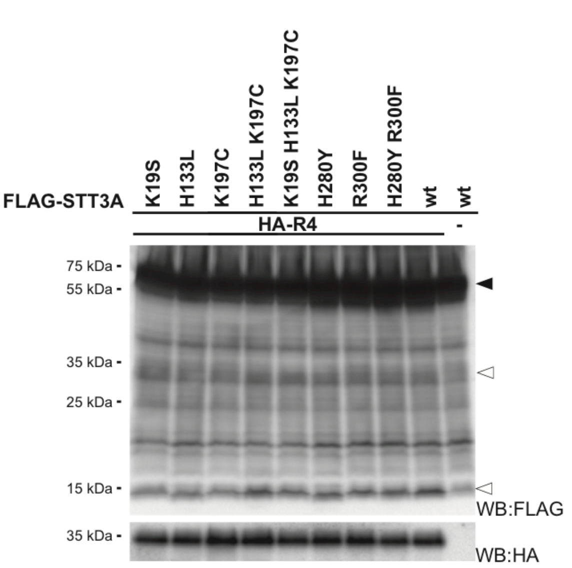


**C**

```
TMD1    RPN2 VSNTFTALILSPLLLLFALWI

TMD2    RPN2 FAPSTIIFHLGHAAMLGLMYV

TMD3    RPN2 LNMFQTLKYLAILGSVTFLAG
```

**D**

```
TMD   RPN1          PLLVVAAFYILFFTVIIYV

TMD   DDOST        YYASAFSMMLGLFIFSIVFL

TMD   B3GNT1 CAFYQLLLAALMLVAMLQLLYLSLLSGL
```

**E**

```
HEc GlpG_P09391        HFSLMHILFNLLWWWYLGGAVEKR

Hs  RHBDL1_O75783      HVGLEQLGFNALLQLMIGVPLEMV
Hs  RHBDL2_Q9NX52      HAGVQHILGNLCMQLVLGIPLEMV
Hs  RHBDL3_P58872      HAGIEHLGLNVVLQLLVGVPLEMV
Mm  RHBDL1_Q8VC82      HVGLEQLGFNALLQLMIGVPLEMV
Mm  RHBDL2_A2AGA4      HAGVQHIVGNLLMQIVLGIPLEMV
Mm  RHBDL3_P58873      HAGVEQLGLNVALQLLVGVPLEMV
Dr  RHBDL1_E7FCU6      HVGLEQLGFNALLQLMIGVPLEMV
Dr  RHBDL2_Q7ZUN9      HAGVEHIMGNLLMQLLLGIPLELV
Dr  RHBDL3_Q566N3      HAGIEHLGLNMAMQLLVGVPLEMV
consensus A-type       *.*::::  *  :*:::*:***:*      A-type

Hs  RHBDL4_Q8TEB9      HADDWHLYFNMASMLWKGINLERR
Mm  RHBDL4_Q8BHC7      HGDDWHLYFNMVSMLWKGVKLERR
Dr  RHBDL4_Q568J3      HVDDMHLYFNMALLLWKGIRLEKK
consensus B-type       * ** ******. :****:.**::      B-type

consensus total        * .  ::  *      *  :*
```

**F**

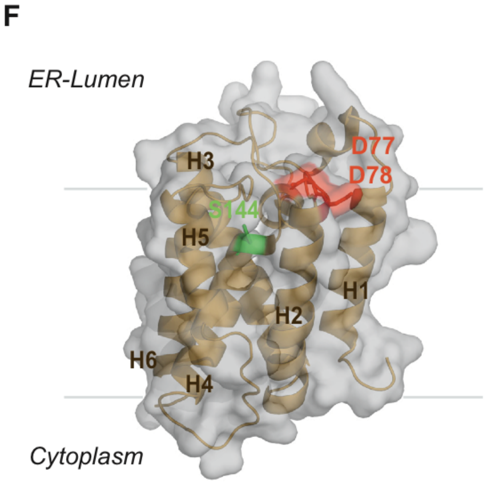


**G**

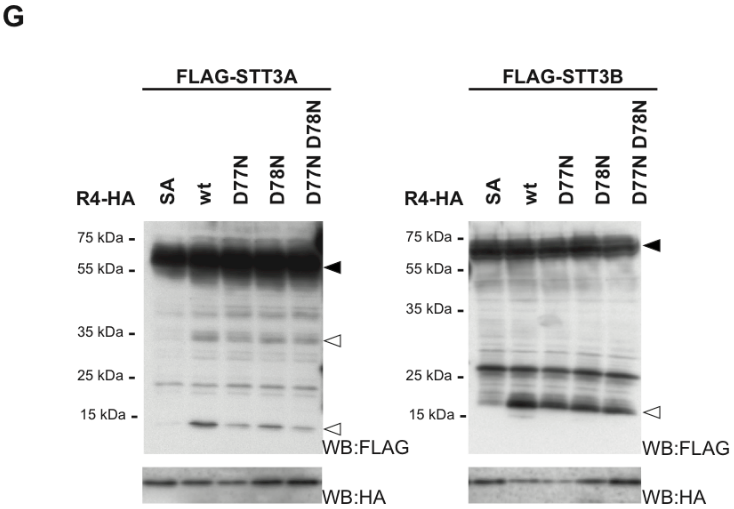

**Fig. S4. STT3A and STT3B show differently charged TM Domains.** (A) Alignment of STT3A and STT3B TM domains (TMD). Positive charged residues are highlighted in blue. Positive charges in STT3A TMDs absent in STT3B are highlighted in yellow. (B) FLAG-STT3A wt or FLAG-STT3A with positive charged TMD residues replaced as indicated with the respective residues of STT3B, have been transfected either with empty vector (-) or HA-tagged RHBDL4 (HA-R4) and cells have been treated with MG132 (2 μM). WB, western blot. (C) RPN2 TMDs. Positive charged residues are highlighted in blue. (D) TMDs of RPN1, DDOST and B4GAT1. (E) Multiple-sequence alignment of TM segment H2 from *E. coli* (Ec) GlpG and secretory pathway rhomboids from human (*H. sapiens*, Hs), mouse (*M. musculus*, Mm) and *D. rerio* (Dr) shows two conserved aspartate residues D77/D78 (highlighted in red) that are missing in the other RHBDLs. Sequence consensus of the A-class and B-class secretase rhomboids according to (Lemberg and Freeman, 2007) are shown. Functionally important residues forming the putative oxyanion hole important for catalysis are highlighted in green (Strisovsky, 2016). (F) Homology model of RHBDL4 reveals position of the putative negative charged substrate docking site. Position of the active site serine-144 (S144) in the active site groove and aspartate-77 (D77) and aspartate-78 (D78) in the putative gating TM segment H2 (Cho et al., 2019) are indicated. (G) Hek293T cells were transfected with FLAG-STT3A or FLAG-STT3B (filled triangle) together with either RHBDL4 SA, wt or RHBDL4 variants where negative charged aspartate residues have been replaced by uncharged asparagines (D77N, D78N, D77N/D78N). RHBDL4 generated cleavage fragments at 13 kDa and 35 kDa for STT3A and at 16 kDa for STT3B are highlighted by open triangles. Cells have been treated with MG132 (2 μM). See Fig. 5D for quantification.

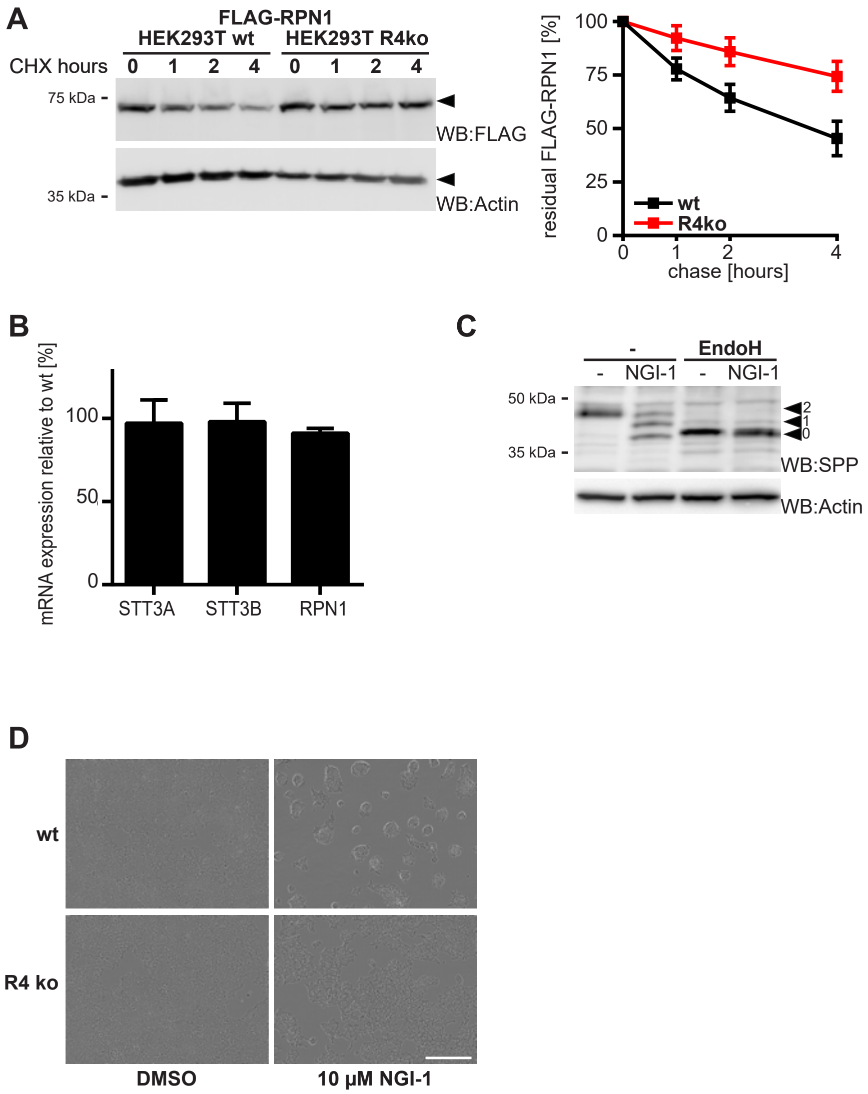


**Fig. S5. Functional characterization of RHBDL4 knockout cells.** (A) Hek293T cells, either wt or RHBDL4 knock-out (R4ko) transiently expressing FLAG tagged RPN1 (FLAG-RPN1) were subjected to cycloheximide (CHX) chase and analyzed by western blotting (WB). Actin serves as loading control. Right panel: Quantification of RPN1-FLAG expression level relative to respective timepoint 0 (means ± SEM, n=3). (B) Transcriptional levels of STT3A, STT3B and RPN1 in HEK293T R4ko cells relative to Hek293T wt cells normalized to the arithmetic mean of the housekeeping genes β-2 microglobulin and TATA-binding protein. (C) Glycosylation status of SPP in Hek293T cells after treatment with DMSO or 10 µM NGI-1 for 24 h analyzed by WB. NGI-1-treatment results in appearance of mono and unglycosylated SPP. Treatment with Endo H removes all N-linked glycans. (D) Images of the cells treated with OST inhibitor NGI-1 or vehicle control (DMSO) depicted in Fig. 6D. 10-fold magnification of cells 60 hours after start of the growth curve. Scale bar: 400 µm.

**Table S1. Complete list of proteins identified by RHBDL4 substrate trapping MS/MS.**
See separate file.

**Table S2. Substrate Candidates identified by RHBDL4 substrate trapping MS/MS.**
Table S1 selected for the following criteria: quantified in both replicates, >= 2 unique peptides, GO term “integral to membrane” and “endoplasmic reticulum”, mean Heavy over Light ratio <=2.0. Number of transmembrane domains has been added manually.

**Table S3. Proteins identified by protein pilot search to determine sequons for N-linked glycosylation – Protein Summary.**
See separate file.

**Table S4. Proteins identified by protein pilot search to determine sequons for N-linked glycosylation – Peptide Summary.**
See separate file.

**Table S5. Proteins identified by protein pilot search to determine sequons for N-linked glycosylation – Disctinct Peptide Summary.**
See separate file.

**Table S6. Ratios of site-specific N-linked occupancy for partially occupied sequons.**
See separate file.
